# Acute inhibition of centriolar satellite function and positioning reveals their functions at the primary cilium

**DOI:** 10.1101/2020.02.10.941658

**Authors:** Ozge Z. Aydin, Sevket Onur Taflan, Can Gurkaslar, Elif Nur Firat-Karalar

**Affiliations:** Department of Molecular Biology and Genetics, Koc University, Istanbul, Turkey

## Abstract

Centriolar satellites are dynamic, membrane-less granules composed of over 200 proteins. They store, modify, and traffic centrosome and primary cilium proteins, and help to regulate both the biogenesis and some functions of centrosomes and cilium. In most cell types, satellites cluster around the perinuclear centrosome, but their integrity and cellular distribution are dynamically remodeled in response to different stimuli, such as cell cycle cues. Dissecting the specific and temporal functions and mechanisms of satellites and how these are influenced by their cellular positioning and dynamics has been challenging using genetic approaches, particularly in ciliated and proliferating cells. To address this, we developed a chemical-based trafficking assay to rapidly and efficiently redistribute satellites to either the cell periphery or center, and fuse them into stable clusters in a temporally-controlled way. Induced satellite clustering at either the periphery or center resulted in antagonistic changes in the pericentrosomal levels of a subset of proteins, revealing a direct role for their positioning in protein targeting and sequestration. Systematic analysis of the interactome of peripheral satellite clusters revealed enrichment of proteins implicated in cilium biogenesis and mitosis. Importantly, inducible peripheral satellite targeting in ciliated cells revealed a function for satellites not just for efficient cilium assembly, but also in the maintenance of steady-state cilia and in cilia disassembly by regulating the structural integrity of the ciliary axoneme. Finally, although perturbing satellite distribution and dynamics inhibited their mitotic dissolution, it did not cause mitotic defects. Collectively, our results for the first time showed a direct link between satellite functions and their pericentrosomal clustering, and provided a new tool for probing temporal satellite functions in different contexts.

## Introduction

The mammalian centrosome/cilium complex consists of the centrosome, cilium, and centriolar satellites, which together regulate polarity, signaling, proliferation and motility in cells and thereby development and homeostasis in organisms. Centriolar satellites (hereafter satellites) are 70-100 nm membrane-less electron-dense granules that localize and move around centrosomes and cilia [1]. Satellite assembly is scaffolded by the large coiled-coil protein Pericentriolar Material 1 (PCM1), which physically interacts with many known and putative centrosome proteins [2, 3]. Satellite resident proteins function in a wide range of cellular processes including cilium assembly, ciliary transport, centriole duplication, mitotic regulation, and microtubule dynamics and organization, and include proteins mutated in developmental and neuronal disorders [4]. Accordingly, acute or constitutive loss of satellites through depletion or deletion of PCM1 resulted in defects in cilium assembly, ciliary signaling, epithelial cell organization and autophagy [4–7]. Satellites mediate their functions in part by regulating the cellular abundance or centrosomal and ciliary targeting of specific proteins [5–8]. These results, together with their microtubule-mediated active transport, led to the current model, which defines satellites as trafficking modules and/or storage sites for their protein residents.

Although satellites are ubiquitous structures in vertebrate cells, their number and distribution changes in a context-dependent way. For example, in most cell types, satellites are predominently clustered around the centrosome, and to a lesser extent scattered throughout the cytoplasm [9]. However, in specialized cell types, their distribution varies from clustering at the nuclear envelope in myotubes and at the apical side of polarized epithelial cells, to being scattered throughout the cell body in neurons [9–11]. In addition to their cell type-specific distribution, satellite properties can also be influenced by extracellular stimuli. For example, genotoxic stress causes satellites to disperse throughout the cytoplasm, and during mitosis, satellites dissolve and subsequently reassemble upon mitotic exit [12, 13]. The mitotic dissolution of PCM1 is regulated by a dual tyrosine kinase 3 (DYRK3)-dependent mechanism and is a line of evidence for liquid-like behaviour of satellites. Consistent with such behaviour, satellite granules were shown to undergo fusion and fission events with each other and the centrosome in epithelial cells [14]. These context-dependent variations in satellite properties suggest that their cellular distribution and remodeling have important implications for their functions, although the mechanisms remain unknown.

There are two key unknowns that pertains to our understanding of satellite functions as discrete protein complexes. First, despite its highly regulate nature, whether, and if so, how satellite distribution contributes to their functions is not known. Second, previous approaches that used depletion or deletion of PCM1 to study satellite functions were limited in uncovering their temporal functions such as the ones during cilium maintenance and mitosis. Addressing these questions is an essential step in defining their relationship with centrosomes and cilia, and thereby the inter-organelle communication within the vertebrate centrosome/cilium complex. To this end, we used chemical manipulation to efficiently and rapidly target satellites to the microtubule plus ends at the cell periphery or microtubule minus ends at the cell center. Satellites targeted to these locations formed stable assemblies, and we showed that satellite proximity to centrosomes and cilia is required for their functions in centrosomal protein targeting and cilium assembly, maintenance and disassembly.

## Results and Discussion

### Development and validation of the inducible satellite trafficking assay

Satellites cluster around the centrosome most cell types, suggesting that their proximity to the centrosomes is important for their functions. To uncover temporal functions of satellites and to elucidate how their pericentrosomal clustering contributes to their functions, we developed an inducible satellite trafficking assay to target satellites away from the centrosomes to the cell periphery and determined the molecular and cellular consequences (Fig. 1A). To correlate the associated phenotypes with satellite proximity to the centrosomes, we in parallel targeted satellites to the cell center (Fig. 1A). Our approach makes use of the inducible dimerization of the FK506 binding protein 12 (FKBP) and FKBP12-rapamycin-binding (FRB) domain of mTOR by rapamycin or its cell-permeable analog rapalog [15, 16]. Due to the long half-life of rapamycin interaction with FKBP and FRB (about 17.5 hours), the dimerization is essentially irreversible [17]. This FKBP-FRB-based rapamycin-inducible heterodimerization approach was previously used to spatiotemporally manipulate specific cellular structures and catalytic activities such as the removal of IFT20 and IFT74 from cilia and sequestration at the mitochondria [18], ciliary targeting of CCP5 tubulin deglutamylase [19], and recruitment of molecular motors to endosomes, peroxisomes and the nucleus [20–23].

**Figure 1.**
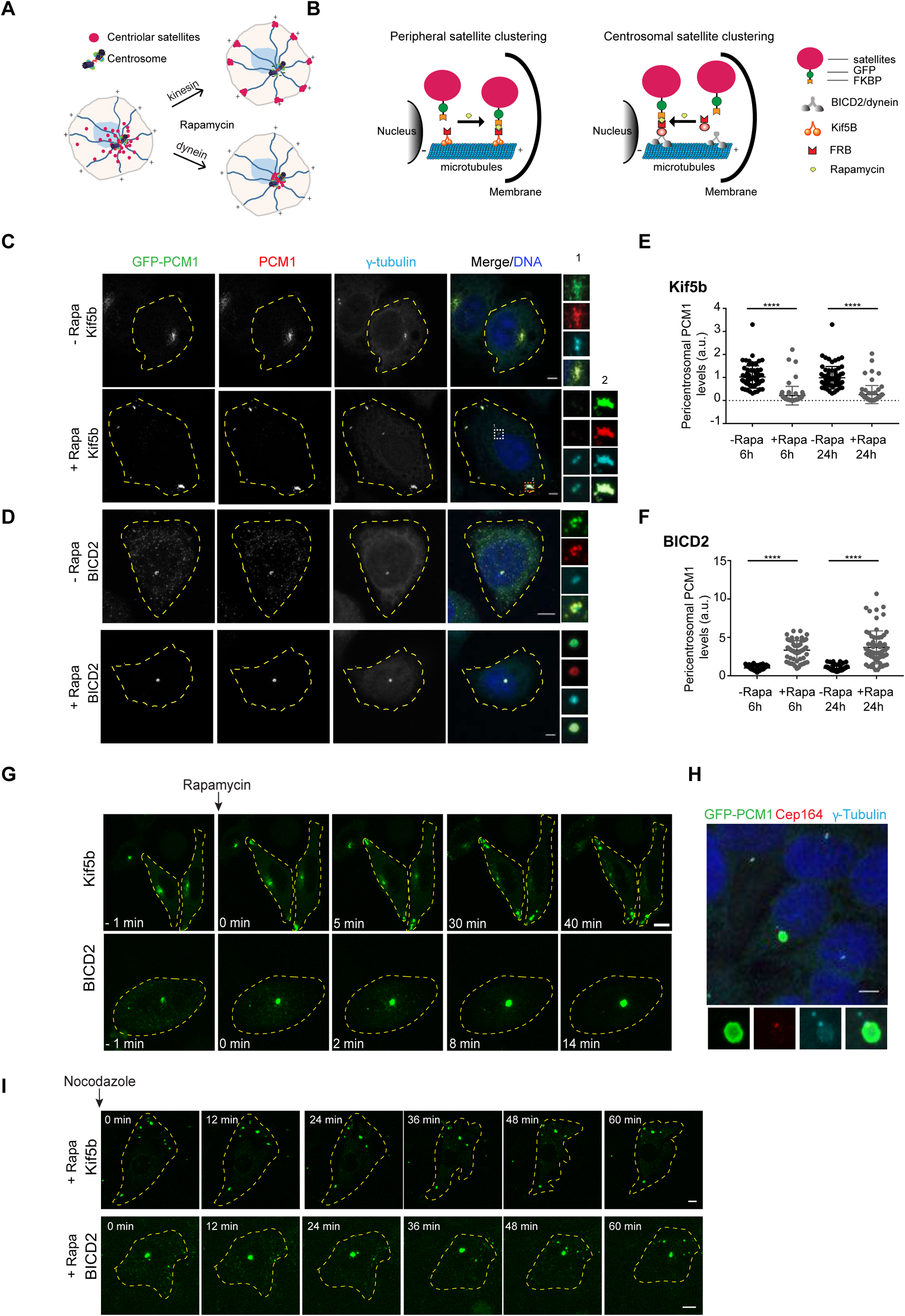
Inducible dimerization of PCM1 with molecular motor domains targets satellites to the cell periphery or center. Development and validation of the inducible satellite trafficking assay. **(A)** Representation of satellite redistribution to cell periphery or center upon inducible dimerization of the satellite scaffolding protein PCM1 with the constitutively active plus and minus end-directed molecular motor proteins. **(B)** Design of the inducible satellite-trafficking assay. PCM1 was tagged with GFP at the N-terminus and FKBP at the C-terminus. Co-expression of GFP-PCM1-FKBP with HA-Kif5b (1-269 a.a.)-FRB targets satellites to the cell periphery where microtubule plus ends are concentrated upon rapamycin addition. Co-expression of GFP-PCM1-FKBP with HA-BICD2 (1-198 a.a.)-FRB targets satellites to cell center where microtubule minus ends are concentrated upon rapamycin addition. **(C, D)** Representative images for satellite distribution in control and rapamycin-treated cells. Cells co-expressing GFP-PCM1-FKBP with HA-Kif5b-FRB or HA-BICD2-FRB were treated with rapamycin for 1h. 24 h after treatment with rapamycin, cells were fixed and stained with antibodies against GFP, PCM1 and g-tubulin. DNA was stained with DAPI. Cell edges are outlined. Scale bar, 5 μm. **(E, F)** Endogenous PCM1 exhibits the same localization pattern as GFP-PCM1-FKBP. Pericentrosomal PCM1 levels was quantified at 6 h and 24 h after rapamycin treatment of **(E)** Kif5b-expressing or **(F)** BICD2-expressing cells. Average mean of pericentrosomal levels in control cells were normalized to 1. n=25 cells per experiment. Data represent mean value from two experiments per condition, ± SD (****p<0.0001). **(G)** Dynamics of satellite redistribution to the cell periphery or center upon rapamycin addition. Cells co-expressing GFP-PCM1-FKBP with HA-Kif5b-FRB or HA-BICD2-FRB were imaged before and after rapamycin addition using time lapse microscopy. Imaging was performed more frequently (every 2 min) in BICD2-expressing cells relative to Kif5b-expressing cells (every 5 min). Representative still frames from time-lapse experiments were shown. Cell edges are outlined. Time, minutes after rapamycin addition. Scale bar, 5 μm. **(H)** Satellites concentrate around mother centriole in a ring-like pattern in cells co-expressing GFP-PCM1-FKBP with HA-BICD2-FRB after rapamycin treatment. Cells were stained for GFP, Cep164 and gamma-tubulin. DNA was stained with DAPI. Cell edges are outlined. Scale bar, 5 μm. **(I)** Dynamics of peripheral and centrosomal satellite clusters upon microtubule depolymerization. Cells with peripheral or centrosomal satellite clustering were treated with nocodazole for 1 h and imaged every 3 min by time lapse microscopy. Representative still frames from time-lapse experiments were shown. Cell edges are outlined. Scale bar, 5 μm.

We adapted this assay for satellites by co-expressing the FKBP-fusion of satellites as the cargo of interest with the FRB-fusion of constitutively active plus-end and minus-end directed motor domains (Fig. 1B). We tagged the satellite-scaffolding protein PCM1 with GFP at the N-terminus for visualization of satellites, and with FKBP at the C-terminus for inducible dimerization with the motor domains (Fig. 1B). Analogous to GFP-PCM1 and endogenous PCM1, GFP-PCM1-FKBP localized to satellites in transfected human cervical carcinoma (HeLa) cells without altering their distribution (Fig. S1A). To target satellites to the microtubule plus ends at the cell periphery, we induced their recruitment to the FRB fusion of constitutively active kinesin-1 HA-Kif5b (1-269 a.a.) motor domain (hereafter HA-Kif5b-FRB or Kif5b), which lacks the tail region required for interaction with endogenous cargoes and thus can only bind to satellites via the FKBP-FRB dimerization (Fig. 1B) [22]. HA-Kif5b-FRB localized to the cytosol and did not affect the distribution of satellites (Fig. S1B). To target satellites to the microtubule minus ends at the cell center, we induced their recruitment to the FRB fusion of the constitutively active N terminus of dynein/dynactin cargo adaptor bicaudal D homolog 2 BICD2 (1-198 a.a.) (hereafter HA-BICD2-FRB or BICD2), which interacts with dynein and dynactin but lacks the cargo-binding domain (Fig. 1B) [22]. HA-BICD2-FRB localized diffusely throughout the cytosol in low-expressing cells and formed granules in high-expressing cells (Fig. S1B). In contrast to HA-Kif5b, we noticed that HA-BICD2 expression by itself resulted in satellite dispersal throughout the cytosol (Fig. S1B), which is consistent with its previous characterization as a dominant-negative mutant that impairs dynein-dynactin function [24–26]. Therefore, we did not use BICD2-based centrosomal satellite clustering as a way to assay satellite functions in ciliated and proliferating cells.

To test the feasibility of the inducible satellite trafficking assay, we first determined the localization of satellites in transfected HeLa cells before and after rapamycin addition by live imaging and quantitative immunofluorescence. In cells co-expressing GFP-PCM1-FKBP and HA-Kif5b-FRB, satellites had their typical pericentrosomal clustering pattern in the absence of rapamycin (Fig. 1C, S1C). Satellite localization ranged from clustering around the centrosomes to scattering throughout the cytosol in cells co-expressing GFP-PCM1-FKBP and HA-BICD2 depending on ectopic BICD2 expression levels (Fig. 1D, S1D). Upon rapamycin addition to cells expressing HA-Kif5b-FRB and GFP-PCM1-FKBP, satellites were targeted to the cell periphery where the majority of microtubule plus ends localize (Fig. 1C, 1G, S1C, MovieS1). In contrast, in cells expressing HA-BICD2-FRB and GFP-PCM1-FKBP, rapamycin addition resulted in clustering of satellites at the centrosome (Fig. 1D, 1G, S1D, MovieS2).

Notably, we observed partial distribution of both GFP-PCM1-FKBP and PCM1 to the cell periphery or center upon rapamycin addition in a subset of cells, likely due to insufficient expression of the active motors relative to the satellites (Fig. S1E). The integrity and organization of the microtubule network remained unaltered in both cases upon rapamycin-induced redistribution of satellites (Fig. S1F).

Importantly, localization of endogenous PCM1 tightly correlated with the localization of GFP-PCM1-FKBP before and after rapamycin addition (Fig. 1C, 1D), which is in agreement with PCM1 self-dimerization through its N-terminal coiled-coil domains [9]. To determine the targeting efficiency of endogenous PCM1, we quantified its pericentrosomal levels by measuring fluorescence signal intensities within a 3 µm^2^ circle encompassing the centrosome. We found that pericentrosomal levels decreased significantly in Kif5b-expressing (peripherally targeted) cells (6 h: Control: 1 ± 0.07, Rapa: 0.25 ± 0.06, p value <0.0001, 24 h: Control: 1 ± 0.07, Rapa: 0.3 ± 0.06, p value <0.0001) and increased significantly in BICD2-expressing (centrosomally targeted) cells (6 h: Control: 1 ± 0.04, Rapa: 3.0 ± 0.17, p value <0.0001, 24 h: Control: 1 ± 0.05, Rapa: 4.1 ± 0.3, p value <0.0001) at 6 h and 24 h after rapamycin induction (Fig. 1E, 1F). As controls, rapamycin treatment of control cells or cells expressing only GFP-PCM1-FKBP did not perturb satellite distribution in cells, as assessed by staining cells with anti-PCM1 antibody (Fig. S1G). Together, these results validated rapid and efficient redistribution of satellites to the cell periphery or center upon rapamycin-induced dimerization.

In Kif5b-expressing cells, satellites formed large clusters that were heterogeneously distributed at the peripheral cell protrusions, whereas in BICD2-expressing cells, they formed a single cluster at the centrosome (Fig. 1C, 1D). A similar heterogeneous distribution pattern at the periphery was also observed when GFP-PCM1-FKBP was co-expressed with the motor domain of another kinesin Kif17 (Fig. S1H). Staining of BICD2-expressing cells for satellites and the mother centriole protein Cep164 revealed tight clustering of satellites around the mother centriole in a ring-like pattern (Fig. 1H). Given that a subset of centrosomal microtubules are anchored at the subdistal appendages of the mother centrioles, this localization pattern suggests preferential trafficking of satellites along these microtubules [27]. Because satellite targeting to the cell periphery or center was dependent on microtubules, we next examined whether these satellite cluster(s), once they formed, were maintained independently of microtubules or not. The clusters remained mostly intact and did not split into smaller granules upon depolymerization of microtubules by nocodazole treatment (Fig. 1I, Movie S3, S4). Together, these results show that rapamycin-induced dimerization of satellites with molecular motors compromises satellite distribution and size irreversibly, and thus this assay provides a tool to investigate satellite functions in a temporally-controlled manner.

### Satellites exert variable effects on pericentrosomal abundance of their residents

Previous work has shown that the cellular loss of satellites, by acute or chronic depletion of PCM1, alters the centrosomal levels and dynamics of a subset of proteins [4, 5, 14, 28]. Given that satellites interact with and store a wide range of centrosome proteins, their pericentrosomal localization is likely required for efficient protein targeting to the centrosomes. To test this, we co-transfected HeLa cells with the indicated constructs and used quantitative immunofluorescence to measure the pericentrosomal levels of various known satellite proteins. We chose satellite proteins that also localize to different parts of the centrosome (e.g., pericentriolar material, distal appendages) and are implicated in different centrosome/cilium-associated cellular functions (Fig. 2A, 2B). As controls, we quantified the transfected cells in which GFP-PCM1-FKBP localized like endogenous PCM1 and excluded the ones that were impaired for pericentrosomal satellite clustering due to overexpression of molecular motors. For quantifying centrosomal levels in cells with peripheral or centrosomal satellite clustering after rapamycin treatment, we only accounted for the cells that exhibited complete redistribution to the cell center or periphery.

**Figure 2.**
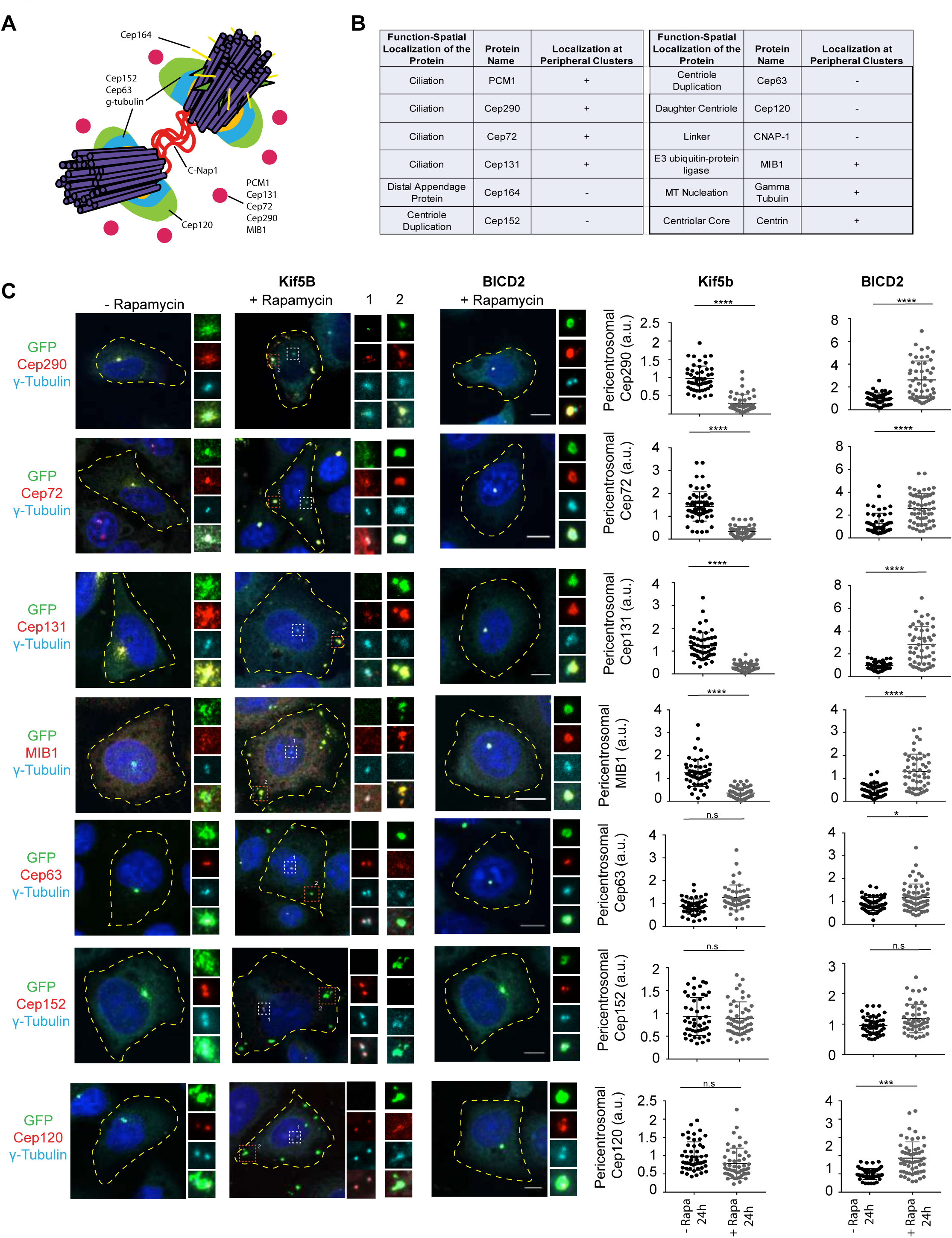
Proper satellite distribution is required for pericentrosomal abundance of satellite residents at varying levels. Effects of satellite mispositioning on pericentrosomal abundance of select satellite residents. **(A)** Spatial localization of the quantified proteins at the centrosome. **(B)** Summary of the results for the changes in the pericentrosomal abundance of the indicated proteins and their associated functions and spatial localizations at the centrosome and cilia. “Localization at peripheral clusters” represent the group of proteins that concentrate with PCM1 at the periphery upon rapamycin addition to Kif5b-expressing cells. **(C)** HeLa cells co-expressing GFP-PCM1-FKBP with HA-Kif5b-FRB or HA-BICD2-FRB were treated with rapamycin for 1 h followed by fixation at 24 h. Cells that were not treated with rapamycin were processed in parallel as controls. Cells were stained with anti-GFP to identify cells with complete redistribution to the cell periphery or center, anti-g-tubulin to mark the centrosome and antibodies against the indicated proteins. Images represent centrosomes in cells from the same coverslip taken with the same camera settings. DNA was stained by DAPI. Fluorescence intensity at the centrosome was quantified and average means of the levels in control cells were normalized to 1. n=25 cells per experiment. Data represent mean value from two experiments per condition, ± SD (***p<0.001, ****p<0.0001). Cell edges are outlined. Scale bars = 10 μm, all insets show 4X enlarged centrosomes.

The pericentrosomal levels of the ciliogenesis factors, including the transition zone component Cep290, the E3 ubiquitin ligase Mib1, Cep131 and Cep72, increased significantly in centrosomally-targeted BICD2-expressing cells and decreased significantly in peripherally-targeted Kif5b-expressing cells at 6 h and 24 h after rapamycin treatment (Fig. 2C, S2A). We also note that these proteins were recruited to the PCM1-positive clusters at the cell periphery in Kif5b-expressing cells (Fig. 2B, 2C). Pericentrosomal accumulation of other satellite proteins implicated in cilium assembly and centriole duplication including KIAA0753, CCDC14 and OFD1 were affected similarly upon satellite mispositioning (Fig. S2B). In contrast, there was no significant change in the pericentrosomal levels of the centriole duplication factors Cep63 and Cep152, the distal appendage protein Cep164, the centriole component centrin and the centrosomal linker protein C-Nap1 in cells with peripheral satellite clustering (Fig. 2B, S2A). Additionally, except for centrin, these proteins were not recruited to the peripheral aggregates in Kif5b-expressing cells.

We next examined the potential role of satellite distribution in regulating the daughter centriole composition by using the rapamycin-treated BICD2-expressing cells, which leads to satellite accumulation around the mother centriole (Fig. 1I). We chose the daughter centriole protein Cep120 for further study as it was identified in the satellite proteome and 60% of its centrosomal pool was shown to be mobile [3, 29, 30]. We stained HeLa cells co-expressing GFP-PCM1-FKBP and HA-BICD2 before and after rapamycin addition with Cep120, the centrosome marker gamma-tubulin, and the mother centriole marker Cep164 (Fig. S3A). Despite its enrichment in the daughter centriole in control cells, Cep120 was strongly enriched in the mother centriole in BICD2-expressing cells (Fig. 3A). This indicates that satellite positioning regulates the association of Cep120 with the daughter centriole.

**Figure 3.**
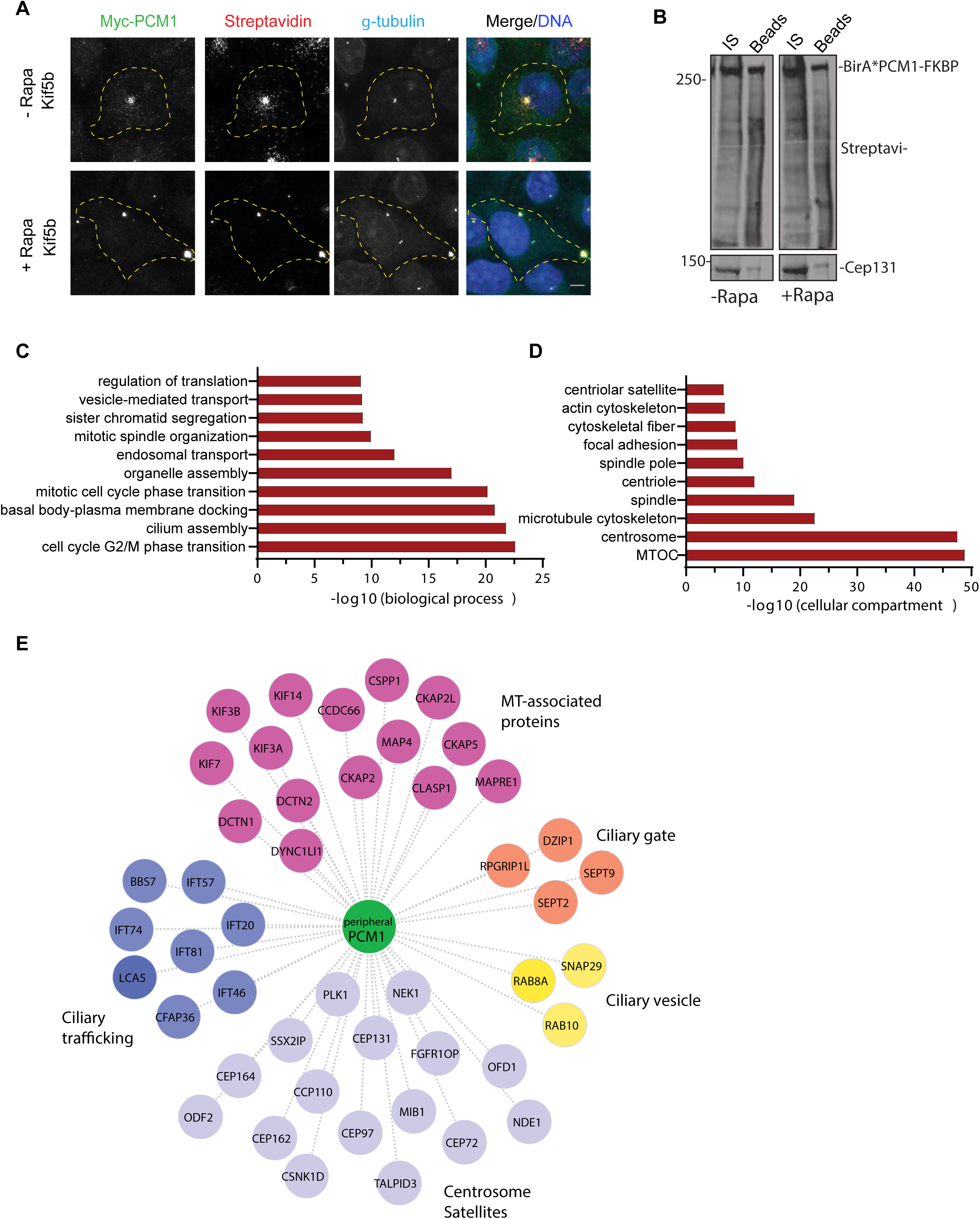
The proximity interactome of PCM1 at the peripheral clusters is enriched for proteins implicated in ciliogenesis and mitosis. The proximity PCM1 interactome of satellites were identified using the BioID approach. **(A)** HEK293T cells were transfected with Myc-BirA*-PCM1-FKBP and induced for peripheral targeting of satellites with rapamycin treatment for 1 h. Cells that were not treated with rapamycin were used as a control. After 18 h biotin incubation, cells were fixed and stained for Myc-BirA*-PCM1-FKBP expression with anti-myc, biotinylated proteins with streptavidin and centrosomes with g-tubulin. DNA was stained with DAPI. Scale bar = 10 μm, cell edges are outlined. **(B)** Biotinylated proteins from cell lysates from cells expressing Myc-BirA*-PCM1-FKBP (-rapamycin or +rapamycin) were pulled down with streptavidin chromatography and samples were analyzed by SDS-PAGE and western blotting with streptavidin to detect biotinylated proteins and with anti-Cep131 (positive control). IS: initial sample used for streptavidin pulldowns, Beads: Eluted proteins. **(C, D)** GO-enrichment analysis of the proximity interactors of PCM1 after rapamycin treatment based on their **(C)** biological processes and **(D)** cellular compartment. The X-axis represents the log transformed p-value (Fishers exact test) of GO terms. **(E)** The cilium-associated proteins in the interactome of peripheral satellites were determined based on GO categories and previous work and different ciliogenesis functional modules plotted in the “peripheral PCM1 interaction network using Cytoscape.

Satellite interactome is highly enriched in microtubule-associated proteins including the key regulators of microtubule nucleation and dynamics such as gamma-tubulin and the components of the HAUS/augmin complex [2, 3]. While the centrosomal gamma-tubulin levels did not change in cells with peripheral satellite clustering, gamma-tubulin concentrated at the peripheral satellite clusters (Fig. S3B). To test whether recruitment of gamma-tubulin to these clusters results in microtubule nucleation, we performed microtubule regrowth experiments in IMCD3 cells stably expressing GFP-PCM1-FKBP and HA-Kif5b. After inducing peripheral accumulation of satellites by rapamycin, microtubules were depolymerized by nocodazole treatment. Following nocodazole washout, cells were fixed and stained for alpha-tubulin at the indicated time points after nocodazole washout (Fig. S3C). Despite prominent microtubule nucleation at the centrosomes, microtubule nucleation was not initiated at the peripheral gama-tubulin-positive clusters. Thus, the concentration of gamma-tubulin at the peripheral satellites does not assign them MTOC activity.

### The majority of the satellite interactome remained unaltered upon peripheral targeting

To determine whether, and if so, how the composition of satellites upon their peripheral targeting, we took a systematic approach. To this end, we applied the BioID proximity labeling approach and identified the PCM1 proximity interactome before and after rapamycin treatment in Kif5b-expressing cells. PCM1-FKBP was fused to myc-BirA* at the N terminus and co-expressed with HA-Kif5b in HEK293T cells. Myc-BirA*-PCM1-FKBP localized to and induced biotinylation at the satellites, as assessed by staining for myc, streptavidin and gamma-tubulin (Fig. 3A). As expected, localized biotinylation was observed at the pericentrosomal granules in control cells and at the peripheral satellite clusters in rapamycin-treated cells (Fig. 3A). Given that satellites are biotinylated at the periphery away from the centrosomes, their proteome will likely exclude the contaminating interactions of PCM1 with the centrosome due to their close proximity.

For mass spectrometry experiments, HEK293T cells were co-transfected with HA-Kif5b and Myc-BirA*-PCM1-FKBP, treated with rapamycin 24 h post-transfection for 1 h and incubated with biotin for 18 h. Cells that were not treated with rapamycin and cells expressing myc-BirA* only were used as controls. Efficient streptavidin pulldown of myc-BirA*-PCM1-fKBP and biotinylated proteins were confirmed by western blotting (Fig. 3B). Biotinylated proteins from two technical and two experimental replicates for each condition were identified by mass spectrometry and analyzed by significance analysis of interactome (SAINT) to identify high-confidence interactions (Table S1).

SAINT analysis identified 541 proximity interactions for PCM1 before rapamycin treatment and 601 interactions after rapamycin treatment (Bayesian false discovery rate (BFDR)<0.01) (Table S2). The control and peripheral PCM1 interactomes shared 476 components, while 65 proteins were specific to peripheral satellites and 125 proteins were specific to control cells (Fig. S4A). The proteins enriched (>1.25 fold relative to control) or depleted (<0.75 fold relative to control) at the peripheral satellites with their associated GO biological processes are shown in Table 2 and Fig. S4. The 71% overlap before and after rapamycin treatment shows that the composition of the PCM1 proximity interactome remains mostly unaltered upon peripheral targeting of satellites. The peripheral PCM1 interactome is significantly enriched in previously identified MTOC, centrosome, satellite and microtubule cytoskeleton components based on Gene Ontology (GO) analysis for “cellular compartment” (Fig. 3D). Of note, we did not identify enrichment for components of microtubule cargoes such as endosomes and lysosomes, which shows that the effects of the mispositioning assay are specific to satellites (Fig. 3D).

GO analysis of the proteins identified in the PCM1 interactome of peripheral satellites for biological processes revealed significant enrichment for processes related to cilium assembly and mitotic progression (Fig. 3C). Given that satellite-less epithelial cells were mainly defective in cilium assembly, we aimed to use this dataset to gain into the specific ciliary processes regulated by satellites [6, 7]. To this end, we determined the PCM1 proximity interactors previously associated with the primary cilium and generated an interaction network by classifying these proteins them based on their functions in the ciliogenesis program (Fig. 3D). Among the highly represented categories are the microtubule-associated proteins previously localized to cilia such as CCDC66, CSPP1 and MAP4 and the centrosome/satellite proteins that regulate cilium biogenesis and function. Additionally, ciliary trafficking complexes IFT-B, kinesin 2 and dynein; proteins implicated in ciliary vesicle formation, septins and transition zone components were also identified in the proteome of peripheral satellite clusters.

Collectively, systematic analysis of the composition peripheral satellites suggest cilium- and mitosis-related cellular processes as potential satellite functions.

### Peripheral centriolar satellite clustering results in defective cilium assembly, maintenance and disassembly

Given our results thus far showed that satellites and associated centrosome proteins can be targeted to the periphery rapidly and efficiently in an inducible way, we employed this assay to investigate temporal satellite functions in ciliated and proliferating cells.

Satellite-less PCM1^-/-^ kidney and retina epithelial cells had significant defects in their ability to assemble primary cilia [6, 7, 31]. We therefore first examined whether pericentrosomal clustering of satellites is also required for cilium assembly. To this end, we generated inner medullary collecting duct (IMCD3) and IMCD3::PCM1^-/-^ cells that stably express GFP-PCM1-FKBP and HA-Kif5b-FRB (hereafter IMCD3^peripheral^ and IMCD3 PCM1 KO^peripheral^ cells). Expression of GFP-PCM1-FKBP restored the ciliogenesis defect of IMCD3 PCM1 KO cells, confirming that this fusion protein is fully functional (Fig. S5A). Rapamycin induction resulted in localization of satellites at the cell periphery in both cell lines (Fig. 4A, S5B). To investigate the consequences of perturbing satellite proximity to the centrosome on cilium assembly, IMCD3^peripheral^ and IMCD3 PCM1 KO^peripheral^ cells were treated with rapamycin for 1 h, serum-starved for 48 h, and the percentage of ciliated cells was quantified by staining cells for Arl13b, a marker for the ciliary membrane, and acetylated tubulin, a marker for the ciliary axoneme (Fig. 4B). While 70.9% ± 3.3 of control cells were ciliated, only 52.8% ± 0.9 of IMCD3 PCM1 KO^peripheral^ cells were ciliated (p<0.01) (Fig. 4C). Similarly, only 77.35% ± 1.65% IMCD^peripheral^ cells ciliated relative to 54.4% ± 1.4 control ciliating population (p<0.01) (Fig. 4D). As a control for the ciliation experiments, rapamycin treatment had no effect on the percentage of ciliated IMCD3 cells at the concentrations and incubation times used (p=0.87) (Fig. S5C). Given that IMCD3^peripheral^ and IMCD3 KO^peripheral^ cells behaved similarly in ciliogenesis experiments, we used IMCD3^peripheral^ experiments for subsequent experiments at the cilia.

**Figure 4.**
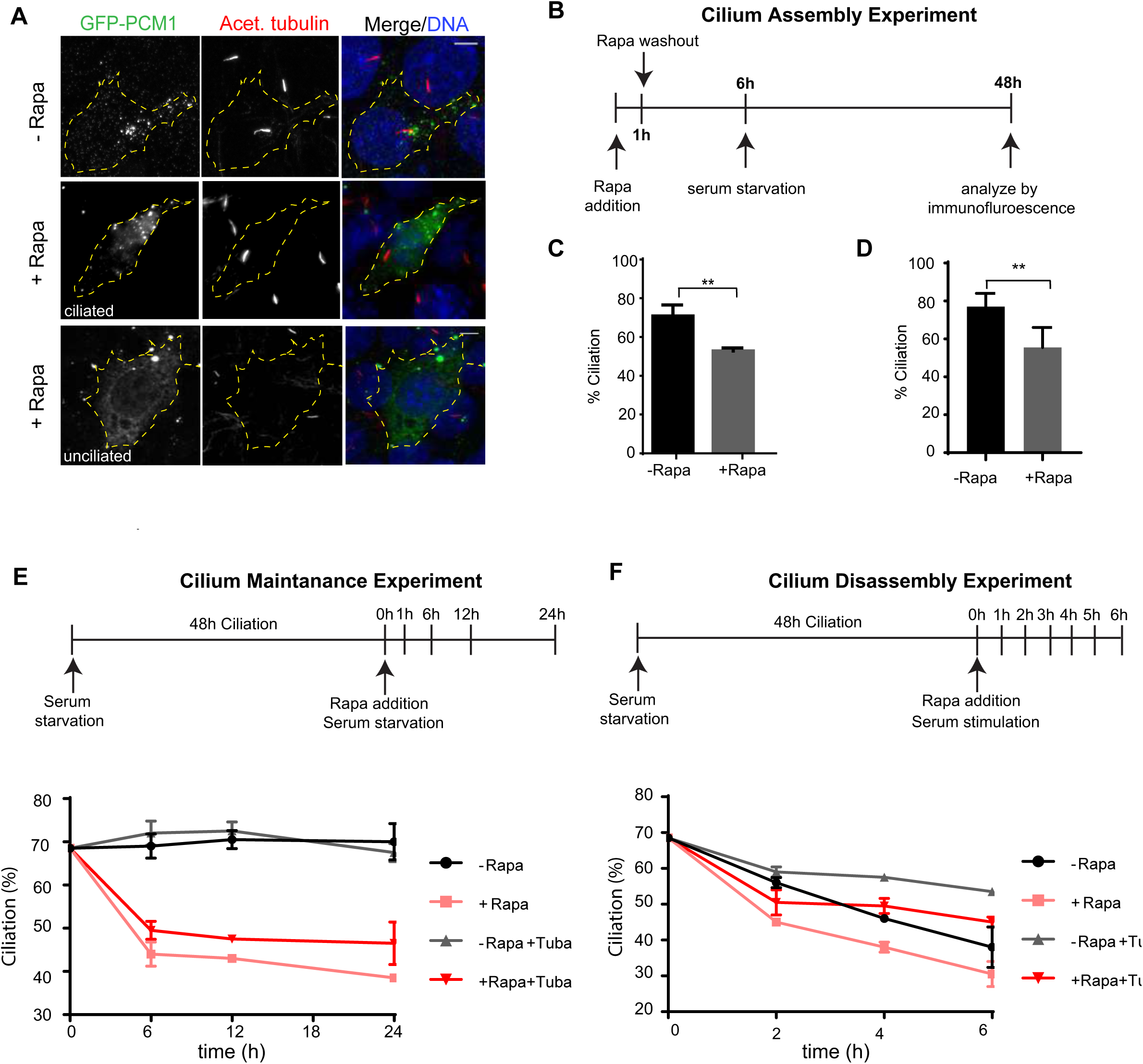
Pericentrosomal satellite clustering is required for efficient cilium assembly, maintenance and disassembly. **(A, B)** Effect of peripheral satellite clustering on cilium assembly. Control and rapamycin-treated IMCD3 PCM1 KO^peripheral^ cells stably expressing GFP-PCM1-FKBP and HA-Kif5b-FRB were serum-starved for 48 h and percentage of ciliated cells was determined by staining for acetylated-tubulin in cells with complete satellite redistribution to the periphery as assessed by GFP staining. Representative images of ciliated and unciliated cells with peripheral satellite clustering relative to ciliated control cells. DNA was stained with DAPI. Scale bar, 10 μm **(C, D)** Quantification of ciliogenesis experiments in **(C)** IMCD3 PCM1 KO^peripheral^ and **(D)** IMCD3^peripheral^ cells. Results shown are the mean of two independent experiments±SD (=50 cells/experiment, **p<0.01). **(E)** Effect of peripheral satellite clustering on cilium maintenance. Cells were serum-starved for 48 h, treated with rapamycin for 1 h and percentage of ciliated cells was determined over 24 h by staining for acetylated-tubulin. Cells that were not treated with rapamycin were used as a control. Same experiments were also performed in the presence of 2 μM tubacin. Data represent mean value from two experiments per condition. **(F)** Effect of peripheral satellite clustering on cilium disassembly. Cells were serum-starved for 48 h, treated with rapamycin for 1 h, induced by serum-stimulation and percentage of ciliated cells was determined over 6 h by staining for acetylated-tubulin. Cells that were not treated with rapamycin were used as a control. Same experiments were also performed in the presence of 2 μM tubacin. Data represent mean value from two experiments per condition.

The inducible nature of the satellite trafficking assay enabled us to study the temporal function of satellites in ciliated cells during primary cilium maintenance and disassembly, which could not have been tested in satellite-less PCM1^-/-^ cells [1, 2]. To study the function of satellites in regulation of steady-state cilia stability, cells were serum starved for 48 h, treated with rapamycin for 1 h, and the percentage of cilia was quantified in a time-course manner after rapamycin washout (Fig. 4E). As expected, the percentage of ciliation did not change over 24 h in control cells (Fig. 4E, Fig. S5D).

However, induction of peripheral satellite clustering resulted in a 22% decrease in the percentage of ciliated cells at 6 h after rapamycin treatment (control: 69% ± 0.5%, Rapa:47.6% ± 3.3, p<0.0001). The fraction of ciliated cells continued to decrease over time until only 38% ± 1% of cells retained primary cilia after 24 h of rapamycin treatment, compared to the stable 69.5% ± 0.5% of control cells (p< 0.0001) (Fig. 4E, Fig. S5D). Rapamycin treatment had no effect on maintenance of control cells (Fig. S5C). Next, we examined whether satellite mispositioning can affect cilium disassembly after serum re-stimulation. To this end, cells were serum starved for 48 h, treated with rapamycin for 1 h, and the percentage of cilia was quantified over a 6h serum-stimulation time course (Fig. 4F, Fig. S5E). In control cells, the percentage of ciliation decreased from 68.5% ± 0.5 to 56% ± 1 at 2 h, 46% ± 0 at 4 h and to 38% ± 4.5 at 6 h (Fig. 4F, Fig. S5E). In rapamycin-treated cells, there was a rapid and significantly greater decrease in the fraction of ciliated cells at 2 h, from 68.5% ± 0.5 to 45% ± 0, which corresponds to the first wave of cilium disassembly [32, 33] (Fig. 4F, Fig. S5E).

After 6 h post serum addition, only 30.5% ± 2.5 of cells with peripheral satellite localization had cilia, compared to 38% ± 4 of control cells (p=0.15) (Fig. 4F, Fig. S5E). The enhanced deciliation and disassembly phenotypes upon peripheral satellite clustering identified satellites as regulators of cilium maintenance and disassembly in addition to their reported functions in assembly.

The maintenance and disassembly of cilium requires modifications of the axonemal tubulins [34]. A major event that promotes cilium disassembly is the phosphorylation of histone deacetylase 6 (HDAC6) by Aurora A kinase and subsequent deacetylation of the modified tubulins of the ciliary axoneme and cortactin [35, 36]. To test whether satellites regulate cilium maintenance and disassembly in a HDAC6-dependent manner, we performed cilium maintenance and disassembly experiments in control and rapamycin-treated cells using tubacin as a potent and selective inhibitor of HDAC6 deacetylase activity [48]. Analogous to control cells, tubacin treatment rescued the enhanced cilium disassembly phenotype of rapamycin-treated cells at 4 and 6 h (4 h: Rapa: 38% ± 4.5%, tubacin+Rapa: 49.5 ± 1.5% (p<0.01); 6h: Rapa: 30.5% ± 2.5%, tubacin+rapa: 45% ± 1%, p<0.01), but did not rescue the cilium maintenance defects at 6,12 or 24 h.(p>0.05) (Fig. 4D, 4E). These results show differential requirements for regulation of cilium maintenance and disassembly, and suggest that satellites function in cilium disassembly by regulating the structural integrity of the ciliary axoneme upstream of HDAC6.

### Satellites are not required for proper mitotic progression and cytokinesis

Satellite integrity and localization is dynamically modulated during cell division and the functional significance of these changes remains poorly understood [5, 37]. The enrichment of key regulators of microtubule nucleation and dynamics, and mitosis such as NUMA, Plk1 and CDK1 in the satellite proteome suggests functions for satellites during mitosis [2, 3]. Unexpectedly, satellite-less IMCD3 and RPE1 PCM1 KO cells did not exhibit defects in cell proliferation, cell cycle progression and mitotic times [1]. The lack of mitotic functions in these cells could be due to the activation of compensatory mechanisms that rescues defects upon constitutive loss of satellites. To investigate whether, and if so, how satellites contribute to mitosis, we assayed the mitotic behavior of IMCD3 PCM1 KO^peripheral^ cells after rapamycin induction of peripheral satellite clustering. IMCD3 PCM1 KO^peripheral^ cells were treated with rapamycin for 1 h followed by time-lapse imaging over 16 h. In control cells, as expected, satellites dissolved during mitosis, which was accompanied by an increase in the soluble cytoplasmic pool of PCM1 (Fig. 5A). In rapamycin-treated cells, the satellite clusters at the periphery lost their peripheral association. Notably, the clusters did not dissolve during mitosis and preferentially accumulated at the cytokinetic bridge (Fig. 5A). Similar mitotic behavior for centriolar satellites were observed in HeLa cells co-expressing GFP-PCM1-FKBP and HA-Kif5b after rapamycin treatment (Fig. 5B).

**Figure 5.**
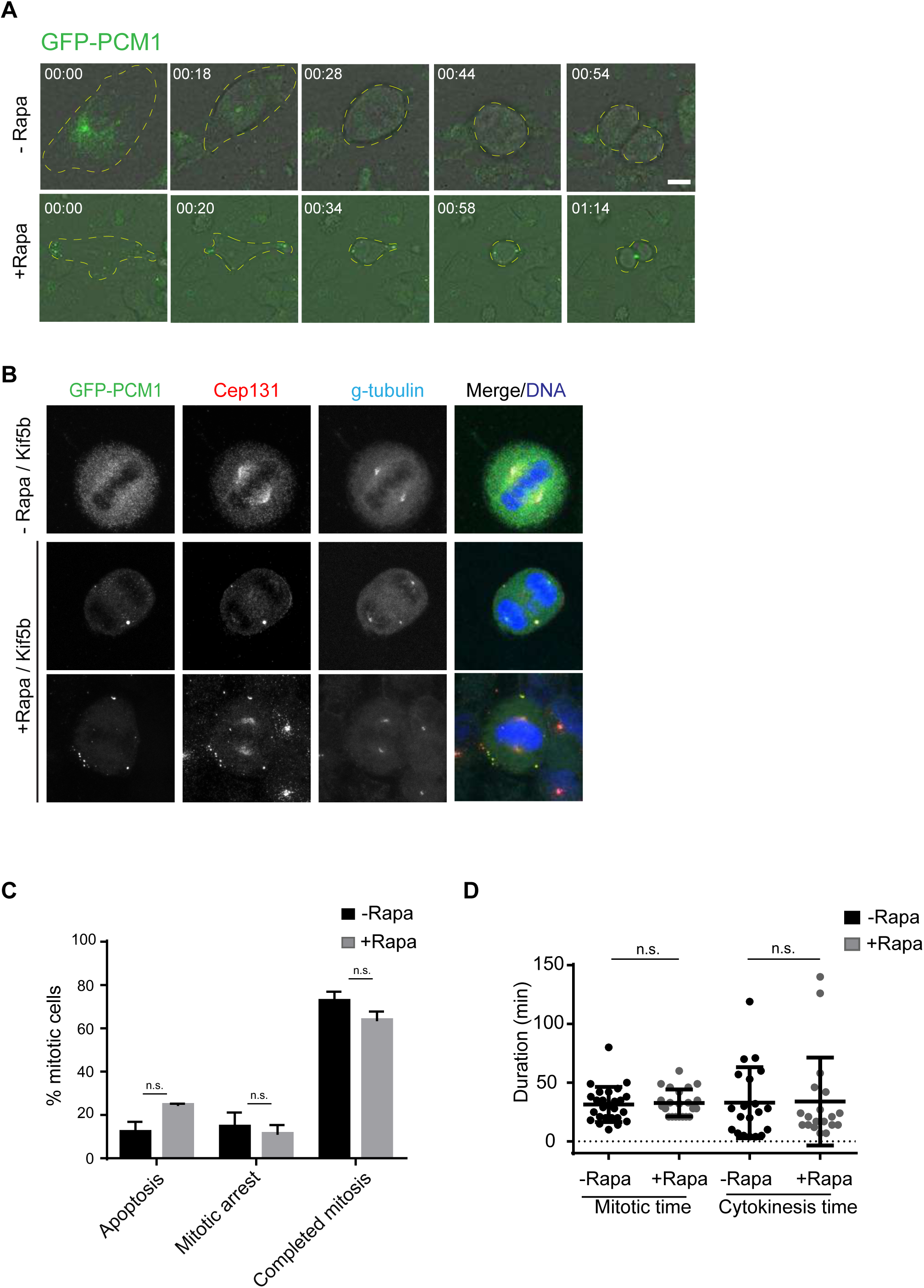
Pericentrosomal satellite clustering is not required for mitotic progression. **(A)** Effect of satellite mispositioning on mitotic dynamics of satellites and mitotic progression. IMCD3 PCM1 KO^peripheral^ cells were imaged every 4.5 min for 16 h. Satellite clusters did not dissolve during mitosis in rapamycin-treated cells, but dissolved in control cells. Representative brightfield and fluorescence still frames from time-lapse experiments were shown. Scale bar, 10 μm **(B)** Effect of satellite mispositioning on satellite integrity during mitosis. Control and rapamycin-treated HeLa cells were fixed 24 h after rapamycin treatment and stained for GFP, Cep131, g-tubulin and DAPI. **(C)** Effect of satellite mispositioning on mitotic progression. Percentage of control and rapamycin-treated mitotic cells were imaged by the time-lapse imaging and videos were analyzed to classify mitotic cells into categories of cells completed mitosis, cells that underwent apoptosis and cells arrested in mitosis for more than 5 h. n>20 cells per experiment. Data represent mean ± SD of three independent experiments (ns: not significant). **(D)** Effect of satellite mispositioning on mitotic and cytokinesis times. Mitotic time was quantified as the time interval from nuclear envelope breakdown to anaphase onset. Cytokinesis time was quantified as the time form anaphase onset to cleavage. n>20 mitotic cells per experiment. Data represent mean ± SD of two independent experiments (ns: not significant).

To examine the phenotypic consequences of satellite mispositioning and inhibition of their mitotic dissolution, we analyzed the movies before and after rapamycin treatment to quantify the percentage of mitotic cells that 1) completed mitosis, 2) underwent apoptosis, 3) arrested in mitosis for more than 5 h. In cells with peripheral satellite clustering, there was not significant change in the fraction of mitotic cells that arrested in mitosis or underwent apoptosis, as assessed by membrane blebbing and DNA fragmentation (3 independent experiments, p>0.05) (Fig. 5C). Next, we quantified the mitotic and cytokinesis time for the group of control and rapamycin-treated cells that completed mitosis during the time of imaging. Mitotic time was determined as the time from nuclear envelope breakdown to anaphase onset by using mCherry-H2B dynamics or brightfield images as references. While the average mitotic time was 31.4±2.9 min for IMCD3^peripheral^ cells that are not treated with rapamycin (n=26), it was 32.8±2.4 for IMCD3^peripheral^ cells after rapamycin treatment (n=27) (2 independent experiments, p>0.05) (Fig. 5D). Cytokinesis time was quantified as the time frame between anaphase onset and cleavage and it was similar between control and rapamycin-treated cells (-rapamycin: 32,95 ± 6,7 min (n=20), + rapamycin: 48,56 ± 12,94 (n=27)) (Fig. 5D).

Rapamycin treatment by itself did not affect mitotic and cytokinesis times (Fig. S5F). Collectively, these results provide further support that satellites are not required for proper mitotic progression and cytokinesis.

## Discussion

We developed a chemically-inducible satellite trafficking assay that allowed us to rapidly and specifically target satellites to the cell periphery and we used this new tool to study satellite functions in a temporally-controlled way in ciliated and proliferating cells.

Application of this assay in epithelial cells for the first time identified functions for satellites not just for cilium assembly, but also for cilium maintenance and disassembly. Targeted and systematic quantification of the satellite proteome at the cell periphery suggested that defects in the pericentrosomal abundance of proteins implicated in these processes as the mechanism underlying these defects. In addition to probing temporal satellite functions, our results for the first time also revealed a direct link between satellite functions and cellular positioning. Importantly, we note that the satellite-trafficking assay provides a powerful tool for future studies aimed at investigating cell-type specific functions and mechanisms of satellites. Its use in specialized cell types, in particular the ones with non-centrosomal MTOCs such as differentiated muscle cells and polarized epithelial cells, also has the potential to provide new insight into why and how satellite distribution varies in different contexts.

Inducible peripheral satellite targeting in ciliated cells revealed novel functions for satellites as positive regulators of cilium maintenance and disassembly. Cilium disassembly upon serum stimulation in IMCD3 cells occurs predominantly by whole-cilium shedding, which requires HDAC6 activity [32]. Enhanced cilium disassembly defect of rapamycin-induced IMCD3^peripheral^ cells was partially restored by HDAC6 inhibition, suggesting that satellites functions upstream of HDAC6. Because HDAC6 was not identified in the satellite interactome, it is likely that satellites are not direct regulators of HDAC6 activity and that they might take part in cilium disassembly through regulating other disassembly regulators such as PLK1 [2, 3]. Of note, PLK1 was identified in the proteome of peripheral satellite clusters. Notably, tubacin treatment did not rescue the cilium maintenance defect, highlighting differential regulation of cilium maintenance and disassembly through independent and/or overlapping mechanisms. Given that we identified the IFT-B components including IFT20, IFT81, IFT57, IFT46 and IFT74 in the proteome of peripheral satellites in Kif5b-expressing cells, we propose that perturbation of IFT trafficking through sequestration of IFT components at the peripheral satellite clusters might underlie the cilium maintenance defects. Supporting this mechanism, sequestration of IFT20 and IFT74 at the mitochondria using rapamycin-dimerization approach in ciliated fibroblasts resulted in a similar defect in cilium maintenance [18]. Together with previous work that identified regulatory roles for satellites in IFT88 targeting to the basal body and cilia [6], our results indicate the presence of a critical, but poorly understood regulatory relationship between satellites and IFT-B complex. Future studies are required to probe the nature of this relationship.

Depletion or deletion of various centrosome proteins perturbs cellular positioning of satellites either by inducing their dispersal throughout the cytoplasm or tighter concentration at the centrosome [29, 38–40]. The satellite distribution defects have been proposed to underlie the phenotypes associated with loss of these proteins such as defective ciliogenesis. However, these loss-of-function studies remained insufficient in directly testing this proposed link. The results of our study provide direct evidence for the functional significance of proper satellite positioning during primary cilium-associated processes, and thus strengthens the conclusions of the previous work on how perturbed satellite localization might contribute to the function of their residents.

Satellites are proposed to mediate their functions in the context of centrosomes and cilia by trafficking proteins to or away from the centrosome and by sequestering centrosome proteins to limit their recruitment at the centrosome [38]. Because our results demonstrated the functional importance of pericentrosomal satellite clustering, it is compelling to propose a complementary function for microtubules and motors in regulating satellite functions. In addition to active transport of centrosome cargoes along microtubules, we propose that microtubules and motors cooperate to maintain satellite proximity around the centrosome and that this is required for timely and efficient exchange of proteins between centrosomes and satellites. Future studies are required to address several key questions that pertains to these models: What is the precise mechanism that establishes and maintains pericentrosomal satellite clustering? Do satellites exchange material with the centrosome through active transport or simple diffusion? What are the upstream signaling pathways that induce protein release from satellites?

The direct link between cellular satellite positioning and their functions revealed by this study corroborates the notion that functions associated with different cellular compartments are regulated by their distinct spatial distribution within cells [49]. This relationship has been reported for multiple organelles and functions. One example is the nutrient-induced peripheral and starvation-induced perinuclear positioning of lysosomes, which was proposed to coordinate mTOR activity and autophagosome biogenesis during cellular anabolic and catabolic responses [41]. Another example is the requirement for proper mitochondrial positioning during T cell activation to allow sustained local Ca+2 influx and during neuronal differentiation to allow terminal axon branching [42–44].

In summary, our findings provide key insight into the satellite functions as key regulators of primary cilium as well as functional relevance of satellite distribution, dynamics, and size. In addition to addressing these mechanisms through *in vitro* and *in vivo* studies, it is essential in future to identify the molecular players that dictate satellite size, distribution, and material properties in different cell types and in distinct cell states. Development of the recently-developed reversible optogenetic and streptavidin-based tools for spatiotemporal manipulation of organelle positioning in the context of satellites will be required to address some of these questions [49, 50].

## Materials and Methods

### Cell culture, transfection and transduction

Human cervical cancer HeLa and human embryonic kidney HEK293T cells were cultured with DMEM medium (Pan Biotech, Cat. # P04-03590) supplemented with 10% FBS and 1% penicillin-streptomycin. Mouse kidney medulla collecting duct cells IMCD3:Flip-In cells were cultured with Dulbecco’s modified Eagle’s Medium DMEM/F12 50/50 medium (Pan Biotech, Cat. # P04-41250) supplemented with 10% Fetal Bovine Serum (FBS, Life Technologies, Ref. # 10270-106, Lot # 42Q5283K) and 1% penicillin-streptomycin (Gibco, Cat. # 1540-122). All cell lines were authenticated by Multiplex Cell Line Authentication (MCA) and were tested for mycoplasma by MycoAlert Mycoplasma Detection Kit (Lonza).

IMCD3 and HeLa cells were transfected with the plasmids using Lipofectamine 2000 according to the manufacturer’s instructions (Thermo Fisher Scientific). For serum starvation experiments, IMCD3 cells were washed twice with PBS and incubated with DMEM/F12 supplemented with 0.5% FBS for the indicated times. For cilium maintenance experiments, IMCD3 cells were serum starved for 48 h, incubated with rapamycin for 1h and fixed at the indicated times. For serum stimulation experiments, ciliated IMCD3 were incubated with rapamycin for 1 h, then with DMEM/F12 50/50 supplemented with 10% FBS and fixed at the indicated times. For rapamycin induction experiments, cells were treated with cells were treated with 100 nm or 500 nm rapamycin (Milipore. Cat#553210) for 1 h followed by 2X PBS washout. To induce microtubule depolymerization, cells were treated with 10 μg/ml nocodazole (Sigma-Aldrich, Cat. #M1404) or vehicle (dimethyl sulfoxide) for one hour at 37°C. For microtubule regrowth experiments, cells were washed 3X with PBS after microtubule depolymerization, incubated in warm media for the indicated times, fixed and stained.

For HDAC inhibition, cells were treated with 2 μm tubacin (Cayman Chemicals, Cat#13691) for the indicated times. IMCD3 cells stably expressing GFP-PCM1-FKBP and HA-Kif5b were generated by cotransfecting cells with expression vectors followed by selection with 750 ug/ml G418 for two weeks and immunofluorescence-based screens for positive colonies. IMCD3 cells stably expressing mCherry-H2B were generated by infecting cells with mCherry-H2B-expressing lentivirus.

### Plasmids

Full-length cDNA of *Homo sapiens* PCM1 was amplified from peGFP-C1-PCM1 and cloned into peGFP-C1 (Clontech) without stop codon using XhoI and KpnI sites. The cDNA of FKBP was amplified from PEX-RFP-FKBP by PCR and cloned into peGFP-PCM1 using KpnI and BamHI sites. PCM1-FKBP was amplified from the peGFP-C1-PCM1-FKBP by PCR and cloned into pcDNA3.1-myc-BirA* using XhoI and BamHI sites. pC4M-F2E-PEX-RFP-FKBP, PCI-neo-HA-BICD2-N (1-594)-FRB, pCI-neo-HI-Kif5b (1-807)-FRB and p®actin-GFP-FRB-Kif17 (1-546) [21, 22, 45].

### Immunofluorescence and antibodies

Cells were grown on coverslips, washed twice with PBS and fixed in either ice cold methanol at -20°C for 10 minutes or 4% PFA in PBS for indirect immunofluorescence. After rehydration in PBS, cells were blocked with 3% BSA (Capricorn Scientific, Cat. # BSA-1T) in PBS + 0.1% Triton X-100 followed by incubation with primary antibodies in blocking solution for 1 hour at room temperature. Cells were washed three times with PBS and incubated with secondary antibodies and DAPI (Thermo Scientific, cat#D1306) at 1:2000 for 45 minutes at room temperature. Following three washes with PBS, cells were mounted using Mowiol mounting medium containing N-propyl gallate (Sigma-Aldrich). Primary antibodies used for immunofluorescence were mouse anti-acetylated tubulin (clone 6-11B, 32270, Thermo Fischer) at 1:10000, mouse anti gamma tubulin (Sigma, clone GTU-88, T5326) at 1:1000, mouse anti GFP (Thermo Scientific, A-11120, clone 3E6) 1:750, mouse anti alpha tubulin (Sigma, DM1A) at 1:1000, rabbit anti CEP152 (Bethyl, A302-480A) at 1:500, rabbit anti-KIAA0753 (Sigma, HPA023494) at 1:1000,rabbit anti-CCDC14 (Genetex, GTX120754) at 1:1000, rabbit anti-OFD1 (Proteintech, 22851-1-AP) at 1:500, rabbit anti-Cep131 (Proteintech, 25757-1-AP) at 1:1000, mouse anti-centrin 3 (Abnova, H0007070-MO1, clone 3E6) at 1:500, rabbit anti-Cep290 (Abcam, ab84870) at 1:1000, anti-rabbit Cep72 (Bethyl, A301-298A) at 1:1000, rabbit anti-Cep63 (Millipore, 61292) at 1:1000, rabbit anti-MIB1 (Sigma, M5948) at 1:1000, mouse anti-CNAP1 (Santa Cruz, sc 390540) at 1:1000, rabbit anti IFT88 (Proteintech, 13967-1-AP) at 1:250, mouse GT335 (Adipogen, A27791601), rabbit Arl13B (Neuromab, Clone N295B/66) at 1:500. Rabbit anti-PCM1, anti-Cep120 and anti-GFP antibodies were generated and used for immunofluorescence as previously described (Firat-Karalar et al., 2014). Secondary antibodies used for immunofluorescence experiments were AlexaFluor 488-, 568- or 633-coupled (Life Technologies) and they were used at 1:2000. Biotinylated proteins were detected with streptavidin coupled to Alexa Fluor 488 or 594 (1:1000; Life Technologies).

### Microscopy and image analysis

Time lapse live imaging was performed with Leica SP8 confocal microscope equipped with an incubation chamber mounted on an automized stage. For cell cycle experiments, asynchronous cells were plated on LabTek dishes (Thermo Fisher) and imaged at 37°C with 5% CO_2_ with a frequency of 4.5 minutes per frame with 1 µm step size and 12 µm stack size in 512 × 512 pixel format at a specific position using HC PL APO CS2 40x 1.3 NA oil objective. For centrosomal protein level quantifications, images were acquired with Leica DMi8 inverted fluorescent microscope with a stack size of 10 µm and step size of 0.5 µm using HC PL APO CS2 63x 1.4 NA oil objective. Higher resolution images were taken by using HC PL APO CS2 63x 1.4 NA oil objective with Leica SP8 confocal microscope.

Quantitative immunofluorescence for pericentrosomal levels of different proteins was performed by acquiring a z-stack of control and depleted cells using identical gain and exposure settings. The maximum-intensity projections were assembled from z-stacks. The centrosome region for each cell were defined by staining for a centrosomal marker gamma-tubulin. The region of interest that encompassed the centrosome was defined as a circle 3 μm^2^ area centered at the centrosome in each cell. Total pixel intensity of fluorescence within the region of interest was measured using ImageJ (National Institutes of Health, Bethesda, MD) (Schneider et al., 2012). Background subtraction was performed by quantifying fluorescence intensity of a region of equal dimensions in the area proximate to the centrosome. Statistical analysis was done by normalizing these values to their mean. Primary cilium formation was assessed by counting the total number of cells, and the number of cells with primary cilia, as detected by DAPI and acetylated tubulin or Arl13b staining, respectively. Ciliary length was measured using acetylated tubulin or Arl13b as the ciliary length marker. All values were normalized relative to the mean of the control cells (=1).

For functional assays, quantification and analysis were performed only in cells that exhibited complete redistribution of satellites to the cell periphery or center.

Complete redistribution to the cell periphery was defined by lack of centrosomal GFP-PCM1-FKBP signal in the pericentrosomal area within 3 μm^2^ area centered on the centrosome. Complete redistribution to the cell center was defined by the lack of GFP-PCM1-FKBP signal in the cytoplasm beyond the 3 μm^2^ pericentrosomal area. As controls, cells that were not treated with rapamycin in which satellites display their typical distribution pattern were quantified.

### Biotin-streptavidin affinity purification

BioID pulldown experiments were performed as previously described [46]. Briefly, HEK293T cells were transfected with myc-BirA*-PCM1-FKBP or myc-BirA*. 24 h post-transfection, cells were incubated with DMEM supplemented 10% FBS and 50 μM biotin for 18. Cells were lysed in lysis buffer (50 mM Tris, pH 7.4, 500 mM NaCI, 0.4% SDS, 5 mM EDTA, 1 mM DTT, 2% Triton X-100, protease inhibitors) and soluble fractions were incubated overnight at 4° with streptavidin agarose beads (Thermo Scientific). Beads were washed twice in wash buffer 1 (2% SDS in dH2O), once with wash buffer 2 (0.2% deoxycholate, 1% Triton X-100, 500 mM NaCI, 1 mM EDTA, and 50 mM Hepes, pH 7.5), once with wash buffer 3 (250 mM LiCI, 0.5% NP-40, 0.5% deoxycholate, 1% Triton X-100, 500 mM NaCI, 1 mM EDTA and 10 mM Tris, pH 8.1) and twice with wash buffer 4 (50 mM Tris, pH 7.4, and 50 mM NaCI). 10% of the sample was analyzed by Western blotting to assess pulldown and 90% of the sample was analyzed by mass spectrometry.

### Mass spectrometry and data analysis

After on-based tryptic digest of biotinylated proteins, peptides were analyzed by online C18 nanoflow reversed-phase nano liquid chromatography (Dionex Ultimate 3000 RS LC, Thermo Scientific) combined with orbitrap mass spectrometer (Q Exactive Orbitrap, Thermo Scientific). Samples were separated in an in-house packed 75 μm i.d.× 23 cm C18 column (Reprosil-Gold C18, 3 μm, 200 Å, Dr. Maisch) using 75 minute linear gradients from 5-25%, 25-40%, 40-95% acetonitrile in 0.1% formic acid with 300 nL/min flow in 90 minutes total run time. The scan sequence began with an MS1 spectrum (Orbitrap analysis; resolution 70,000; mass range 400–1,500 m/z; automatic gain control (AGC) target 1e6; maximum injection time 32 ms). Up to 15 of the most intense ions per cycle were fragmented and analyzed in the orbitrap with Data Dependent Acquisition (DDA). MS2 analysis consisted of collision-induced dissociation (higher-energy collisional dissociation (HCD)) (resolution 17,500; AGC 1e6; normalized collision energy (NCE) 26; maximum injection time 85 ms). The isolation window for MS/MS was 2.0 m/z. Raw files were processed with Thermo Proteome Discoverer 1.4. Carbamidomethylation of cysteine was used as fixed modification and acetylation (protein N-termini) and oxidation of methionine were used as variable modifications.

Maximal two missed cleavages were allowed for the tryptic peptides. The precursor mass tolerance was set to 10 ppm and both peptide and protein false discovery rates (FDRs) were set to 0.01. The database search was performed against the human Uniprot database (release 2016).

Mass spectrometry data for each sample were derived from two biological and two technical replicates. Spectral counts of identified proteins were used to calculate BFDR and probability of possible protein-protein interaction by SAINTexpress v.3.6.3 with -L option [47]. Proteins were filtered out according to SAINTscore (>0.5) and BFDR (<0.05). Interaction maps were drawn using Cytoscape. GO-enrichment analysis was done using EnrichR. The significantly altered GO categories (biological process and cellular compartment) were selected with a Bonferroni-adjusted cutoff p-value of 0.05.

### Statistical analysis

Statistical results, average and standard deviation values were computed and plotted by using Prism (GraphPad, La Jolla, CA). Two-tailed t-tests and ANOVA were applied to compare the statistical significance of the measurements. Error bars reflect SD. Following key is followed for asterisk placeholders for *p*-values in the figures: \**P* < 0.05, \*\**P* < 0.01, \*\*\**P* < 0.001, **** *P* < 0.0001

## Acknowledgements

We acknowledge all members of Firat-Karalar laboratory and Jennifer Wang from the Stearns laboratory at Stanford University for insightful discussions regarding this work. FKBP and FRB constructs were a kind gift from Lukas Kapitein (Utrecht University). We acknowledge the Koç University Proteomics Facility for mass spectrometry analysis and Altug Kamacioglu for the SAINT analysis of the mass spectrometry data. We acknowledge Life Science Editors for editing assistance. This work was supported by ERC Grant 679140 to ENF, Royal Society Newton Advanced Fellowship to ENF and EMBO Installation Grant to ENF.

## Author Contributions

Conceptualization: ENF, OZA, SOT; Methodology: ENF, OZA, SOT, CG; Validation: ENF, OZA, SOT; Formal analysis: ENF, OZA, SOT; Investigation: OZA, SOT; Resources: ENF; Writing – original draft: ENF, OZA; Writing – review and editing: OZA, SOT; Supervision: ENF, OZA; Project administration: ENF; Funding acquisition: ENF

---

**Figure S1.**
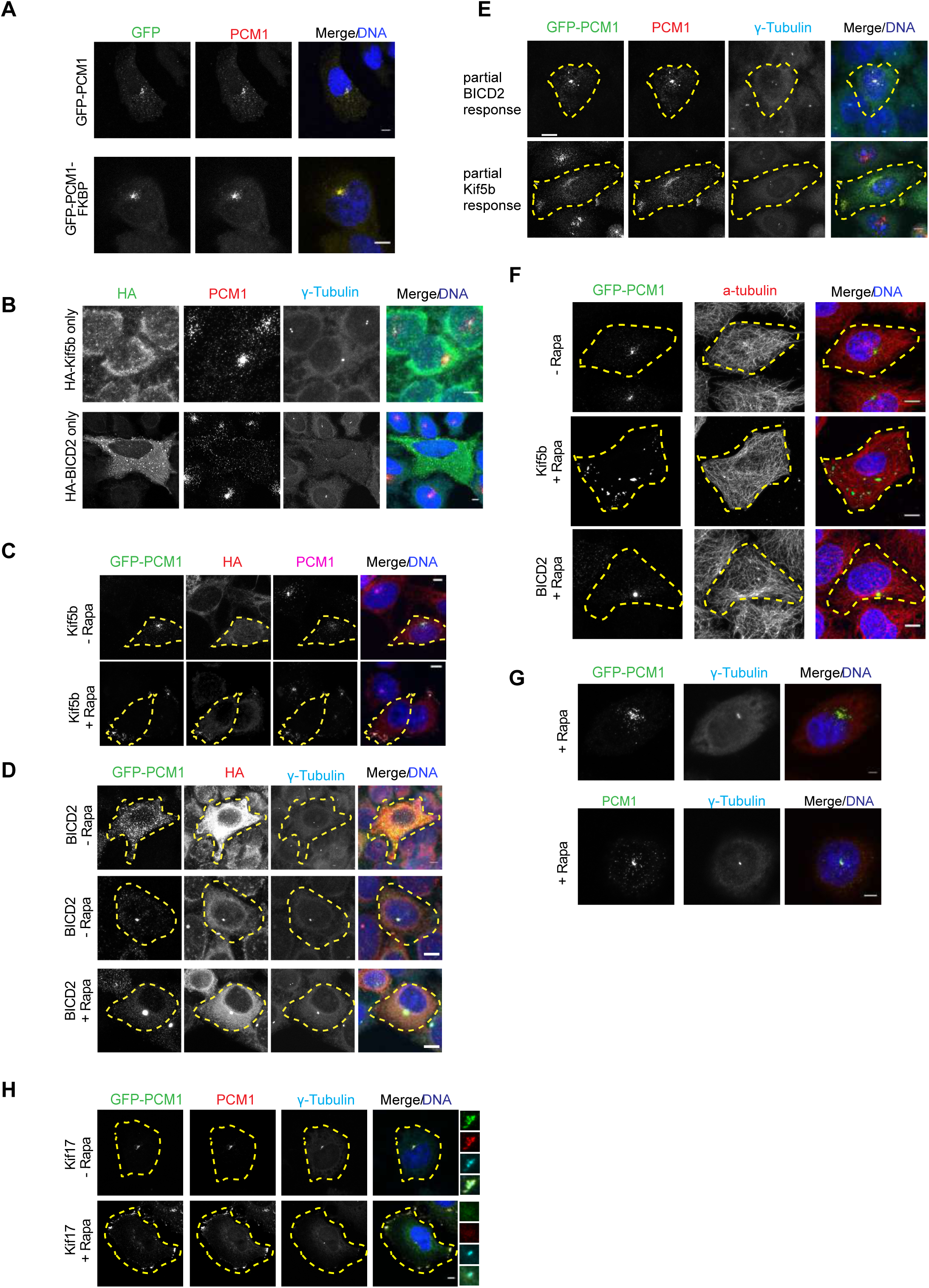
Development and validation of the inducible trafficking assay. **(A)** Localization of GFP-PCM1 and GFP-PCM1-FKBP in cells. HeLa cells were transfected with GFP-PCM1 or GFP-PCM1-FKBP, fixed after 24 h and stained for GFP, PCM1 and DAPI. **(B)** Effects of HA-BICD2 or HA-Kif5b expression on satellite distribution. HeLa cells were transfected with HA-BICD2 or HA-Kif5b, fixed after 24 h and stained for HA, PCM1, g-tubulin and DAPI. **(C, D)** Validation of GFP-PCM1-FKBP and endogenous PCM1 mispositioning upon rapamycin-induced dimerization in cells. Hela cells co-expressing GFP-PCM1-FKBP with **(C)** HA-Kif5b-FRB or **(D)** HA-BICD2- FRB were treated with rapamycin for 1 h, fixed 24 h after transfection and stained for GFP, HA, PCM1 and DAPI. Cells that were not treated with rapamycin were processed in parallel as controls. **(E)** Representation of partial distribution of satellites upon rapamycin induction. Hela cells co-expressing GFP-PCM1-FKBP with HA-Kif5b-FRB or HA-BICD2-FRB were treated with rapamycin for 1 h, fixed 24 h after transfection and stained for GFP, HA, PCM1 and DAPI. Partial distribution was defined by GFP-PCM1- FKBP signal in the pericentrosomal area in Kif5b-expressing cells and in the region excluding therosomal area in BICD2-expressing cells. **(F)** Expression of GFP-PCM1- FKBP with HA-Kif5b-FRB or HA-BICD2-FRB and their redistribution upon rapamycin induction do not perturb the microtubule network. Cells were stained for GFP, alpha- tubulin and DAPI. **(G)** Rapamycin treatment did not perturb satellite distribution in wild-type cells or cells expressing only GFP-PCM1-FKBP. Cells were treated with rapamycin for 1 h, fixed after 24 h and stained for GFP or PCM1, g-tubulin and DAPI. F) Co-expression of GFP-PCM1-FKBP with the constitutively active HA-Kif17 (1-181 a.a.)- FRB targets satellites to the cell periphery where satellite clusters are heterogeneously distributed. Transfected HeLa cells were treated with rapamycin for 1 h, fixed after 24 h and stained for GFP, PCM1, g-tubulin and DAPI. Scale bars = 10 μm, all insets show 4X enlarged centrosomes.

**Figure S2.**
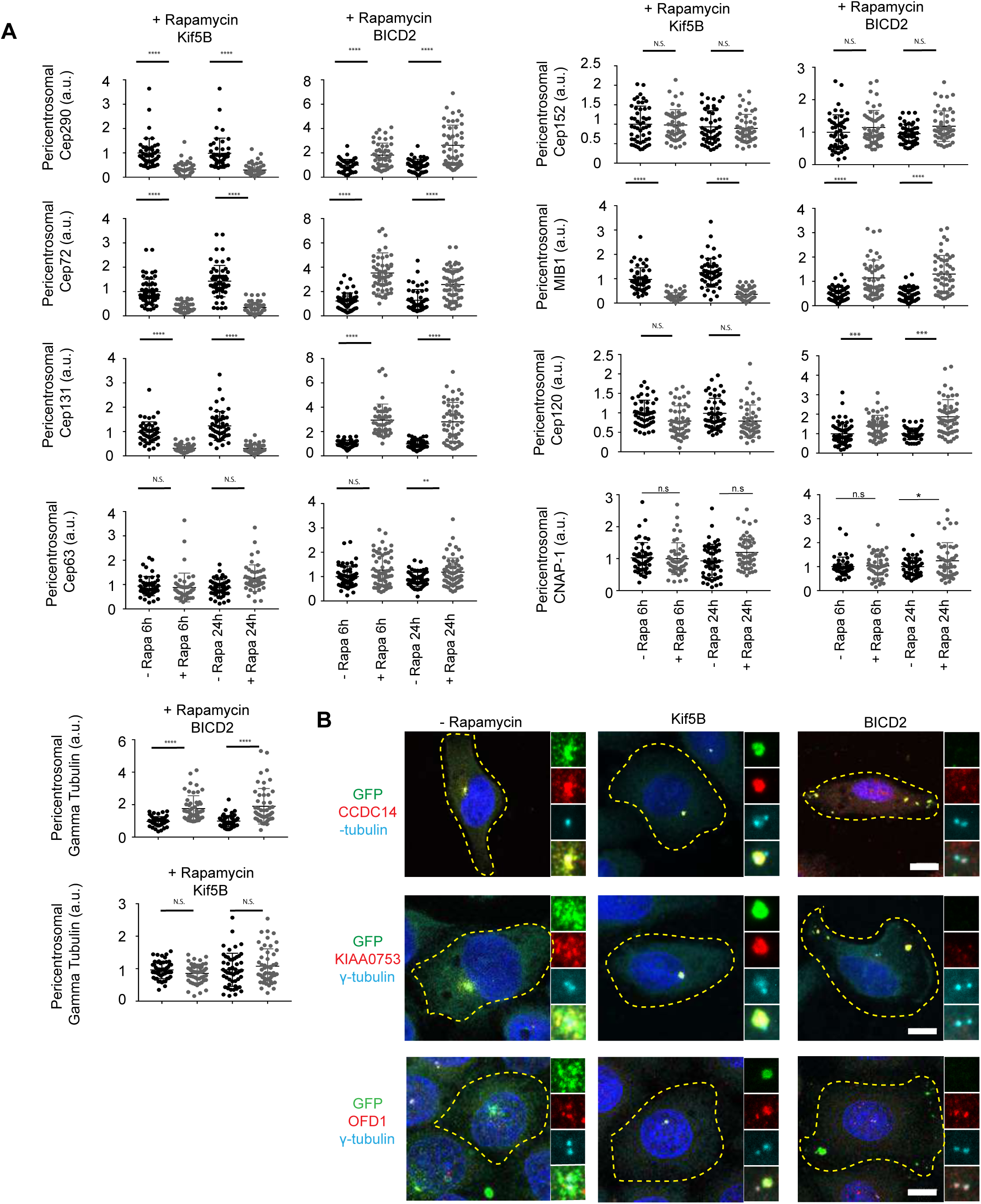
Effects of satellite mispositioning on the pericentrosomal levels of various satellite residents. **(A)** HeLa cells co-expressing GFP-PCM1-FKBP with HA-Kif5b-FRB or HA-BICD2-FRB were treated with rapamycin for 1 h followed by fixation at 6 h and 24 h. Cells that were not treated with rapamycin and exhibited pericentrosomal clustering of GFP-PCM1- FKBP like endogenous PCM1 of wild-type cells were processed in parallel as controls. Cells were stained with antibodies anti-GFP to identify cells with complete redistribution to the cell periphery or center, anti-g-tubulin to mark the centrosome and antibodies against the indicated proteins. Fluorescence intensity at the centrosome was quantified and average means of the levels in control cells were normalized to 1. n=25 cells per experiment. Data represent mean value from two experiments per condition, ± SD (***p<0.001, ****p<0.0001). **(B)** Images represent centrosomes in cells from the same coverslip taken with the same camera settings. DNA was stained by DAPI. Cell edges are outlined. Scale bars = 10 μm, all insets show 4X enlarged centrosomes.

**Figure S3.**
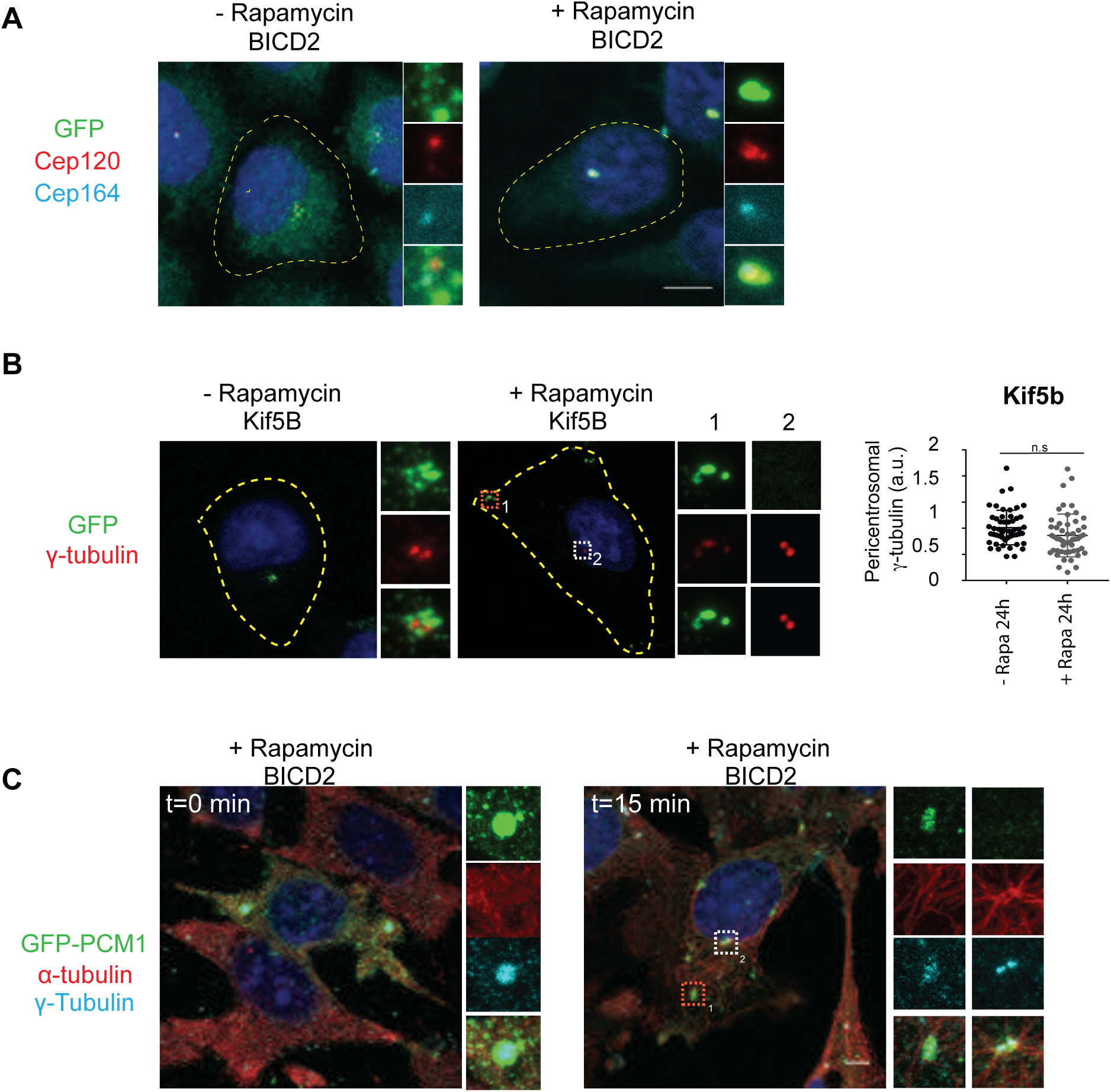
Effects of peripheral satellite clustering on microtubule nucleation and daughter centriole composition. GFP-PCM1-FKBP was co-transfected to HeLa cells with HA-Kif5b-FRB or HA-BICD2- FRB and cells with complete peripheral or centrosomal clustering of satellites after rapamycin induction was assayed for various centrosome functions. **(A)** The daughter centriole protein Cep120 was redistributed to the mother centriole in BICD2-expresing cells with centrosomal satellite accumulation. HeLa cells co-expressing GFP-PCM1- FKBP with HA-BICD2-FRB were treated with rapamycin for 1 h, fixed at 24 h and stained for GFP, Cep120, Cep164 and DAPI. Cells that were not treated with rapamycin were used as a control. **(B)** g-tubulin localization in control cells and in cells with Kif5b- expressing peripheral satellite clustering. HeLa cells co-expressing GFP-PCM1-FKBP with HA-Kif5b-FRB were treated with rapamycin for 1 h, fixed at 24 h and stained for GFP, g-tubulin and DAPI. Images represent centrosomes in cells from the same coverslip taken with the same camera settings. Cells that were not treated with rapamycin were processed in parallel as controls. Fluorescence intensity at the centrosome was quantified and average means of the levels in control cells were normalized to 1. n=25 cells per experiment. Data represent mean value from two experiments per condition, ± SD (n.s. not significant, ****p<0.0001). **(C)** Effect of gamma-tubulin peripheral accumulation on microtubule nucleation. Rapamycin-treated HeLa cells co-expressing GFP-PCM1-FKBP and HA-Kif5b-FRB were treated with DMSO or 10 μg/ml nocodazole for 1 h. After microtubule depolymerization, cells were washed, incubated with media for the indicated times, fixed and stained for GFP, alpha-tubulin, g-tubulin and DAPI. Images represent cells from the same coverslip taken with the same camera settings. Scale bars = 10 μm, all insets show 4X enlarged centrosomes.

**Figure S4.**
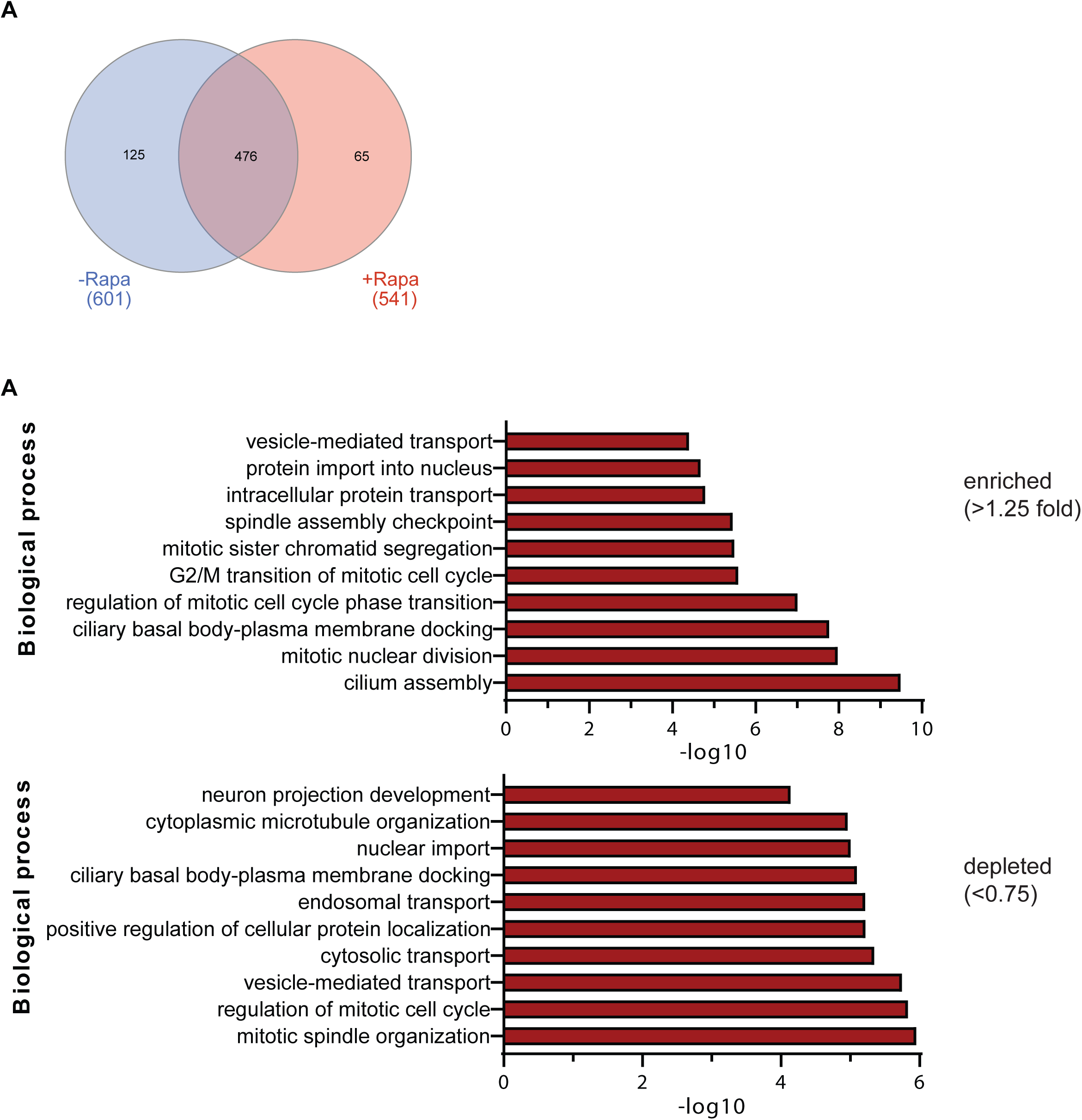
Comparative analysis of the PCM1 proximity interactome of control cells and cells with peripheral satellite targeting. **(A)** The control and peripheral PCM1 interactomes share 476 components, while 65 proteins were specific to peripheral satellites and 125 proteins were specific to control cells. **(B)** GO-enrichment analysis of the proteins enriched (>1.25 fold relative to control) and depleted (<0.75 fold relative to control) in cells with peripheral satellite clustering after rapamycin treatment based on their biological processes. The X-axis represents the log transformed p-value (Fishers exact test) of GO terms.

**Figure S5.**
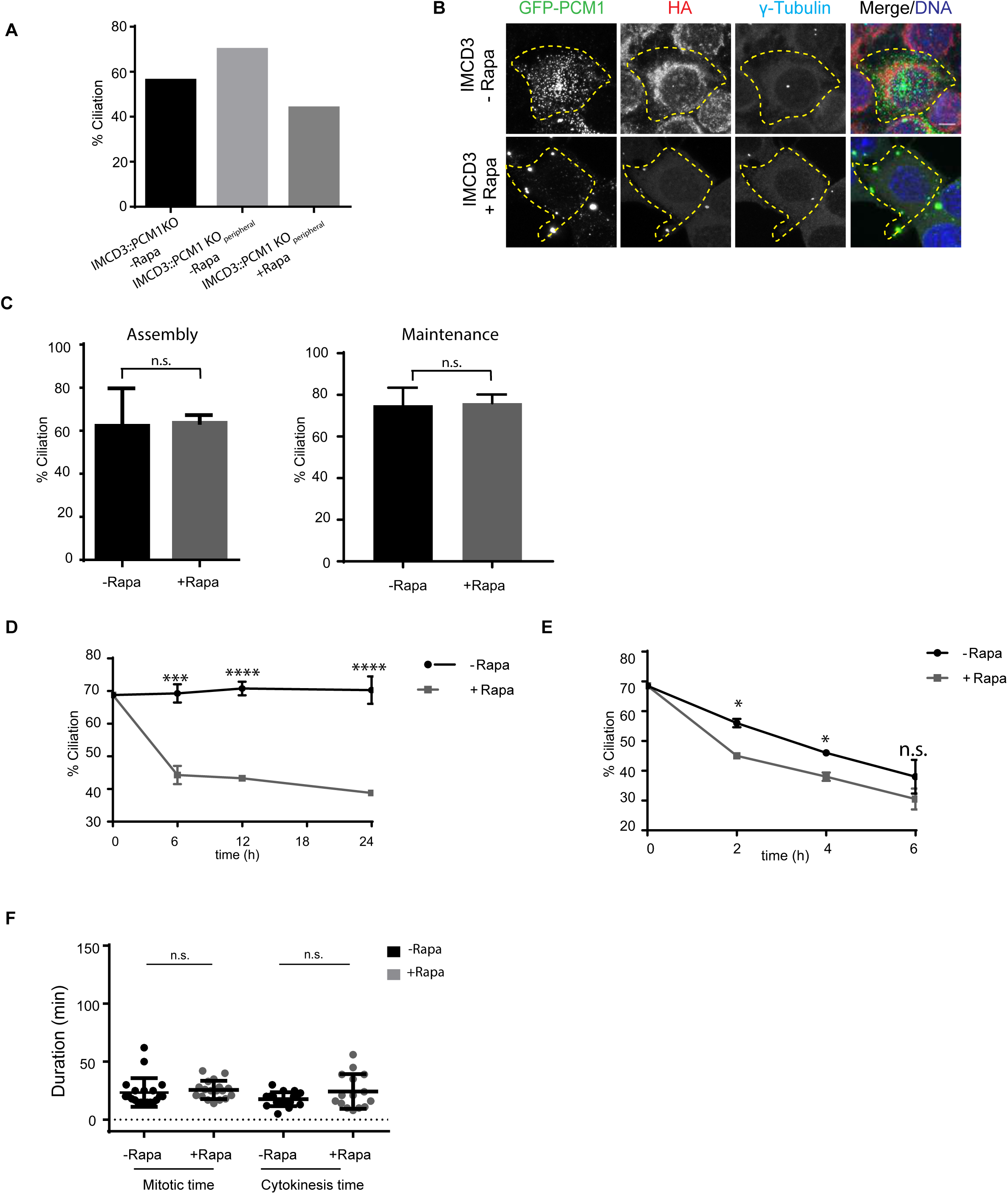
Rapamycin treatment does not interfere with ciliogenesis, cilium maintenance and cilium disassembly. **(A)** GFP-PCM1-FKBP restores ciliogenesis defects of satellite-less IMCD3 PCM1 KO cells. IMCD3 PCM1 KO cells stably expressing GFP-PCM1-FKBP and HA-Kif5b-FRB and IMCD3 PCM1 KO cells were serum-starved for 48 h and percentage of ciliated cells was determined by staining for acetylated-tubulin. **(B)** Validation of IMCD3^perihperal^ cells for expression of HA-Kif5b and GFP-PCM1-FKBP. Control and rapamycin-induced IMCD3^peripheral^ cells were stained for GFP, HA, g-tubulin and DAPI. **(C)** Rapamycin treatment by itself in control IMCD3 cells does not affect the efficiency of ciliogenesis and cilium maintenance. The experiments were performed in IMCD3 cells following the experimental outline for ciliogenesis and maintenance experiment and the fraction of ciliated cells were quantified after 48 h of serum starvation for cilium assembly experiment and 24 h after rapamycin treatment for maintenance experiments. Results shown are the mean of two independent experiments±SD (=200 cells/experiment, n.s: not significant). **(D)** Effect of peripheral satellite clustering on cilium maintenance. Cells were serum-starved for 48 h, treated with rapamycin for 1 h and percentage of ciliated cells was determined over 24 h by staining for acetylated-tubulin. Cells that were not treated with rapamycin were used as a control. Data represent mean value from two experiments per condition, ± SD (***p<0.001, ****p<0.0001). **(E)** Effect of peripheral satellite clustering on cilium disassembly. Cells were serum-starved for 48 h, treated with rapamycin for 1 h, induced by serum-stimulation and percentage of ciliated cells was determined over 6 h by staining for acetylated-tubulin. Data represent mean value from two experiments per condition, ± SD (*p<0.1). **(F)** Rapamycin treatment itself does not affect mitotic time and cytokinesis time of IMCD3 PCM1 KO cells.

**Table S1. Raw mass spectrometry data for Myc-BirA*-PCM1-FKBP with and without rapamycin treatment in Kif5b-expressing cells** Mass spectrometry analysis of proximity interactors of Myc-BirA*-PCM1-FKBP proteins. Proximity interactors from two experimental and two technical replicates for each condition were analyzed by SAINT.

**Table S2. Unique proteins identified in control and rapamycin-treated Myc-BirA*- PCM1-FKBP**

Proteins with BFDR (<0.01) values were defined as part of the PCM1 interactomes.

**Tab 1:** Unique peptides identified in control and rapamycin-treated cells (-Rapa versus +Rapa)

**Tab 2:** Proteins enriched (>1.25 fold relative to control), depleted (<0.75 fold relative to control) and unaltered in rapamycin-treated cells relative to control cells

**MovieS1.** Live imaging of HeLa cells co-expressing GFP-PCM1-FKBP and HA-Kif5b before and after rapamycin treatment

**MovieS2.** Live imaging of HeLa cells co-expressing GFP-PCM1-FKBP and HA-Kif5b before and after rapamycin treatment

**MovieS1.** Live imaging of rapamycin-induced HeLa cells co-expressing GFP-PCM1-FKBP and HA-Kif5b after nocodazole treatment treatment

**MovieS1.** Live imaging of rapamycin-induced HeLa cells co-expressing GFP-PCM1-FKBP and HA-BICD2 after nocodazole treatment treatment

## References

1. Kubo A, Sasaki H, Yuba-Kubo A, Tsukita S, Shiina N. Centriolar satellites: molecular characterization, ATP-dependent movement toward centrioles and possible involvement in ciliogenesis. J Cell Biol. 1999;147(5):969–80. Epub 1999/12/01. PubMed PMID: 10579718.

2. Gheiratmand L, Coyaud E, Gupta GD, Laurent EM, Hasegan M, Prosser SL, et al. Spatial and proteomic profiling reveals centrosome-independent features of centriolar satellites. EMBO J. 2019. Epub 2019/06/05. doi: 10.15252/embj.2018101109. PubMed PMID: 31160316.

3. Quarantotti V, Chen JX, Tischer J, Gonzalez Tejedo C, Papachristou EK, D’Santos CS, et al. Centriolar satellites are acentriolar assemblies of centrosomal proteins. EMBO J. 2019. Epub 2019/06/05. doi: 10.15252/embj.2018101082. PubMed PMID: 31160315.

4. Prosser SL, Pelletier L. Centriolar satellite biogenesis and function in vertebrate cells. J Cell Sci. 2020;133(1). Epub 2020/01/04. doi: 10.1242/jcs.239566. PubMed PMID: 31896603.

5. Dammermann A, Merdes A. Assembly of centrosomal proteins and microtubule organization depends on PCM-1. J Cell Biol. 2002;159(2):255–66. Epub 2002/10/31. doi: 10.1083/jcb.200204023 jcb.200204023 [pii]. PubMed PMID: 12403812.

6. Odabasi E, Gul S, Kavakli IH, Firat-Karalar EN. Centriolar satellites are required for efficient ciliogenesis and ciliary content regulation. EMBO Rep. 2019. Epub 2019/04/27. doi: 10.15252/embr.201947723. PubMed PMID: 31023719.

7. Wang L, Lee K, Malonis R, Sanchez I, Dynlacht BD. Tethering of an E3 ligase by PCM1 regulates the abundance of centrosomal KIAA0586/Talpid3 and promotes ciliogenesis. eLife. 2016;5. doi: 10.7554/eLife.12950. PubMed PMID: 27146717; PubMed Central PMCID: PMC4858382.

8. Joachim J, Razi M, Judith D, Wirth M, Calamita E, Encheva V, et al. Centriolar Satellites Control GABARAP Ubiquitination and GABARAP-Mediated Autophagy. Curr Biol. 2017;27(14):2123–36 e7. doi: 10.1016/j.cub.2017.06.021. PubMed PMID: 28712572; PubMed Central PMCID: PMCPMC5526835.

9. Kubo A, Tsukita S. Non-membranous granular organelle consisting of PCM-1: subcellular distribution and cell-cycle-dependent assembly/disassembly. J Cell Sci. 2003;116(Pt 5):919–28. Epub 2003/02/07. PubMed PMID: 12571289.

10. Srsen V, Fant X, Heald R, Rabouille C, Merdes A. Centrosome proteins form an insoluble perinuclear matrix during muscle cell differentiation. BMC Cell Biol. 2009;10:28. Epub 2009/04/23. doi: 10.1186/1471-2121-10-281471-2121-10-28 [pii]. PubMed PMID: 19383121.

11. Vladar EK, Stearns T. Molecular characterization of centriole assembly in ciliated epithelial cells. J Cell Biol. 2007;178(1):31–42. Epub 2007/07/04. doi: jcb.200703064 [pii] 10.1083/jcb.200703064. PubMed PMID: 17606865.

12. Rai AK, Chen JX, Selbach M, Pelkmans L. Kinase-controlled phase transition of membraneless organelles in mitosis. Nature. 2018;559(7713):211-6. doi: 10.1038/s41586-018-0279-8. PubMed PMID: 29973724.

13. Villumsen BH, Danielsen JR, Povlsen L, Sylvestersen KB, Merdes A, Beli P, et al. A new cellular stress response that triggers centriolar satellite reorganization and ciliogenesis. EMBO J. 2013;32(23):3029–40. Epub 2013/10/15. doi: 10.1038/emboj.2013.223 emboj2013223 [pii]. PubMed PMID: 24121310.

14. Conkar D, Bayraktar H, Firat-Karalar EN. Centrosomal and ciliary targeting of CCDC66 requires cooperative action of centriolar satellites, microtubules and molecular motors. Sci Rep. 2019;9(1):14250. Epub 2019/10/05. doi: 10.1038/s41598-019-50530-4. PubMed PMID: 31582766.

15. Clackson T, Yang W, Rozamus LW, Hatada M, Amara JF, Rollins CT, et al. Redesigning an FKBP-ligand interface to generate chemical dimerizers with novel specificity. Proc Natl Acad Sci U S A. 1998;95(18):10437–42. Epub 1998/09/02. doi: 10.1073/pnas.95.18.10437. PubMed PMID: 9724721; PubMed Central PMCID: PMCPMC27912.

16. Hoogenraad CC, Wulf P, Schiefermeier N, Stepanova T, Galjart N, Small JV, et al. Bicaudal D induces selective dynein-mediated microtubule minus end-directed transport. EMBO J. 2003;22(22):6004–15. Epub 2003/11/12. doi: 10.1093/emboj/cdg592. PubMed PMID: 14609947; PubMed Central PMCID: PMCPMC275447.

17. Hosoi H, Dilling MB, Shikata T, Liu LN, Shu L, Ashmun RA, et al. Rapamycin causes poorly reversible inhibition of mTOR and induces p53-independent apoptosis in human rhabdomyosarcoma cells. Cancer Res. 1999;59(4):886–94. Epub 1999/02/24. PubMed PMID: 10029080.

18. Eguether T, Cordelieres FP, Pazour GJ. Intraflagellar transport is deeply integrated in hedgehog signaling. Mol Biol Cell. 2018;29(10):1178–89. Epub 2018/03/16. doi: 10.1091/mbc.E17-10-0600. PubMed PMID: 29540531; PubMed Central PMCID: PMCPMC5935068.

19. Hong SR, Wang CL, Huang YS, Chang YC, Chang YC, Pusapati GV, et al. Spatiotemporal manipulation of ciliary glutamylation reveals its roles in intraciliary trafficking and Hedgehog signaling. Nat Commun. 2018;9(1):1732. Epub 2018/05/02. doi: 10.1038/s41467-018-03952-z. PubMed PMID: 29712905; PubMed Central PMCID: PMCPMC5928066.

20. Bentley M, Decker H, Luisi J, Banker G. A novel assay reveals preferential binding between Rabs, kinesins, and specific endosomal subpopulations. J Cell Biol. 2015;208(3):273–81. Epub 2015/01/28. doi: 10.1083/jcb.201408056. PubMed PMID: 25624392; PubMed Central PMCID: PMCPMC4315250.

21. Kapitein LC, Schlager MA, Kuijpers M, Wulf PS, van Spronsen M, MacKintosh FC, et al. Mixed microtubules steer dynein-driven cargo transport into dendrites. Curr Biol. 2010;20(4):290–9. Epub 2010/02/09. doi: 10.1016/j.cub.2009.12.052. PubMed PMID: 20137950.

22. Kapitein LC, Schlager MA, van der Zwan WA, Wulf PS, Keijzer N, Hoogenraad CC. Probing intracellular motor protein activity using an inducible cargo trafficking assay. Biophys J. 2010;99(7):2143–52. Epub 2010/10/07. doi: 10.1016/j.bpj.2010.07.055. PubMed PMID: 20923648; PubMed Central PMCID: PMCPMC3042561.

23. Wilson MH, Holzbaur EL. Nesprins anchor kinesin-1 motors to the nucleus to drive nuclear distribution in muscle cells. Development. 2015;142(1):218–28. Epub 2014/12/18. doi: 10.1242/dev.114769. PubMed PMID: 25516977; PubMed Central PMCID: PMCPMC4299143.

24. Guardia CM, De Pace R, Sen A, Saric A, Jarnik M, Kolin DA, et al. Reversible association with motor proteins (RAMP): A streptavidin-based method to manipulate organelle positioning. PLoS Biol. 2019;17(5):e3000279. Epub 2019/05/18. doi: 10.1371/journal.pbio.3000279. PubMed PMID: 31100061; PubMed Central PMCID: PMCPMC6542540.

25. Hoogenraad CC, Akhmanova A, Howell SA, Dortland BR, De Zeeuw CI, Willemsen R, et al. Mammalian Golgi-associated Bicaudal-D2 functions in the dynein-dynactin pathway by interacting with these complexes. EMBO J. 2001;20(15):4041–54. Epub 2001/08/03. doi: 10.1093/emboj/20.15.4041. PubMed PMID: 11483508; PubMed Central PMCID: PMCPMC149157.

26. Teuling E, van Dis V, Wulf PS, Haasdijk ED, Akhmanova A, Hoogenraad CC, et al. A novel mouse model with impaired dynein/dynactin function develops amyotrophic lateral sclerosis (ALS)-like features in motor neurons and improves lifespan in SOD1-ALS mice. Hum Mol Genet. 2008;17(18):2849–62. Epub 2008/06/27. doi: 10.1093/hmg/ddn182. PubMed PMID: 18579581.

27. Piel M, Meyer P, Khodjakov A, Rieder CL, Bornens M. The respective contributions of the mother and daughter centrioles to centrosome activity and behavior in vertebrate cells. J Cell Biol. 2000;149(2):317–30. Epub 2000/04/18. doi: 10.1083/jcb.149.2.317. PubMed PMID: 10769025; PubMed Central PMCID: PMCPMC2175166.

28. Hames RS, Crookes RE, Straatman KR, Merdes A, Hayes MJ, Faragher AJ, et al. Dynamic recruitment of Nek2 kinase to the centrosome involves microtubules, PCM-1, and localized proteasomal degradation. Mol Biol Cell. 2005;16(4):1711-24. Epub 2005/01/22. doi: 10.1091/mbc.e04-08-0688. PubMed PMID: 15659651; PubMed Central PMCID: PMCPMC1073654.

29. Betleja E, Nanjundappa R, Cheng T, Mahjoub MR. A novel Cep120-dependent mechanism inhibits centriole maturation in quiescent cells. eLife. 2018;7. Epub 2018/05/10. doi: 10.7554/eLife.35439. PubMed PMID: 29741480; PubMed Central PMCID: PMCPMC5986273.

30. Mahjoub MR, Xie Z, Stearns T. Cep120 is asymmetrically localized to the daughter centriole and is essential for centriole assembly. J Cell Biol. 2010;191(2):331–46. Epub 2010/10/20. doi: 10.1083/jcb.201003009 jcb.201003009 [pii]. PubMed PMID: 20956381.

31. Stowe TR, Wilkinson CJ, Iqbal A, Stearns T. The centriolar satellite proteins Cep72 and Cep290 interact and are required for recruitment of BBS proteins to the cilium. Mol Biol Cell. 2012;23(17):3322–35. Epub 2012/07/07. doi: 10.1091/mbc.E12-02-0134 mbc.E12-02-0134 [pii]. PubMed PMID: 22767577.

32. Mirvis M, Siemers KA, Nelson WJ, Stearns TP. Primary cilium loss in mammalian cells occurs predominantly by whole-cilium shedding. PLoS Biol. 2019;17(7):e3000381. Epub 2019/07/18. doi: 10.1371/journal.pbio.3000381. PubMed PMID: 31314751; PubMed Central PMCID: PMCPMC6699714.

33. Sanchez I, Dynlacht BD. Cilium assembly and disassembly. Nat Cell Biol. 2016;18(7):711-7. doi: 10.1038/ncb3370. PubMed PMID: 27350441; PubMed Central PMCID: PMCPMC5079433.

34. Mirvis M, Stearns T, James Nelson W. Cilium structure, assembly, and disassembly regulated by the cytoskeleton. Biochem J. 2018;475(14):2329–53. doi: 10.1042/BCJ20170453. PubMed PMID: 30064990; PubMed Central PMCID: PMCPMC6068341.

35. Pugacheva EN, Jablonski SA, Hartman TR, Henske EP, Golemis EA. HEF1- dependent Aurora A activation induces disassembly of the primary cilium. Cell. 2007;129(7):1351–63. Epub 2007/07/03. doi: 10.1016/j.cell.2007.04.035. PubMed PMID: 17604723; PubMed Central PMCID: PMCPMC2504417.

36. Ran J, Yang Y, Li D, Liu M, Zhou J. Deacetylation of alpha-tubulin and cortactin is required for HDAC6 to trigger ciliary disassembly. Sci Rep. 2015;5:12917. Epub 2015/08/08. doi: 10.1038/srep12917. PubMed PMID: 26246421; PubMed Central PMCID: PMCPMC4526867.

37. So C, Seres KB, Steyer AM, Monnich E, Clift D, Pejkovska A, et al. A liquid-like spindle domain promotes acentrosomal spindle assembly in mammalian oocytes. Science. 2019;364(6447). Epub 2019/06/30. doi: 10.1126/science.aat9557. PubMed PMID: 31249032; PubMed Central PMCID: PMCPMC6629549.

38. Hori A, Barnouin K, Snijders AP, Toda T. A non-canonical function of Plk4 in centriolar satellite integrity and ciliogenesis through PCM1 phosphorylation. EMBO Rep. 2016;17(3):326–37. doi: 10.15252/embr.201541432. PubMed PMID: 26755742; PubMed Central PMCID: PMCPMC4772974.

39. Kim J, Krishnaswami SR, Gleeson JG. CEP290 interacts with the centriolar satellite component PCM-1 and is required for Rab8 localization to the primary cilium. Hum Mol Genet. 2008;17(23):3796–805. Epub 2008/09/06. doi: 10.1093/hmg/ddn277 ddn277 [pii]. PubMed PMID: 18772192.

40. Silva E, Betleja E, John E, Spear P, Moresco JJ, Zhang S, et al. Ccdc11 is a novel centriolar satellite protein essential for ciliogenesis and establishment of left-right asymmetry. Mol Biol Cell. 2016;27(1):48–63. Epub 2015/11/06. doi: 10.1091/mbc.E15-07-0474mbc.E15-07-0474 [pii]. PubMed PMID: 26538025.

41. Korolchuk VI, Saiki S, Lichtenberg M, Siddiqi FH, Roberts EA, Imarisio S, et al. Lysosomal positioning coordinates cellular nutrient responses. Nat Cell Biol. 2011;13(4):453–60. Epub 2011/03/12. doi: 10.1038/ncb2204. PubMed PMID: 21394080; PubMed Central PMCID: PMCPMC3071334.

42. Courchet J, Lewis TL, Jr., Lee S, Courchet V, Liou DY, Aizawa S, et al. Terminal axon branching is regulated by the LKB1-NUAK1 kinase pathway via presynaptic mitochondrial capture. Cell. 2013;153(7):1510–25. Epub 2013/06/26. doi: 10.1016/j.cell.2013.05.021. PubMed PMID: 23791179; PubMed Central PMCID: PMCPMC3729210.

43. Korolchuk VI, Rubinsztein DC. Regulation of autophagy by lysosomal positioning. Autophagy. 2011;7(8):927–8. Epub 2011/04/28. doi: 10.4161/auto.7.8.15862. PubMed PMID: 21521941; PubMed Central PMCID: PMCPMC3149695.

44. Schwindling C, Quintana A, Krause E, Hoth M. Mitochondria positioning controls local calcium influx in T cells. J Immunol. 2010;184(1):184–90. Epub 2009/12/02. doi: 10.4049/jimmunol.0902872. PubMed PMID: 19949095.

45. Lipka J, Kapitein LC, Jaworski J, Hoogenraad CC. Microtubule-binding protein doublecortin-like kinase 1 (DCLK1) guides kinesin-3-mediated cargo transport to dendrites. EMBO J. 2016;35(3):302–18. Epub 2016/01/14. doi: 10.15252/embj.201592929. PubMed PMID: 26758546; PubMed Central PMCID: PMCPMC4741305.

46. Conkar D, Culfa E, Odabasi E, Rauniyar N, Yates JR, 3rd, Firat-Karalar EN. Centriolar satellite protein CCDC66 interacts with CEP290 and functions in cilium formation and trafficking. J Cell Sci. 2017. doi: 10.1242/jcs.196832. PubMed PMID: 28235840.

47. Teo G, Liu G, Zhang J, Nesvizhskii AI, Gingras AC, Choi H. SAINTexpress: improvements and additional features in Significance Analysis of INTeractome software. J Proteomics. 2014;100:37–43. Epub 2014/02/12. doi: 10.1016/j.jprot.2013.10.023. PubMed PMID: 24513533; PubMed Central PMCID: PMCPMC4102138.

48. Haggarty SJ, Koeller KM, Wong JC, Grozinger CM, Schreiber SL. Domain-selective small-molecule inhibitor of histone deacetylase 6 (HDAC6)-mediated tubulin deacetylation. Proc Natl Acad Sci U S A. 2003;100(8):4389–94. Epub 2003/04/05. doi: 10.1073/pnas.0430973100. PubMed PMID: 12677000; PubMed Central PMCID: PMCPMC153564.

49. van Bergeijk P, Hoogenraad CC, Kapitein LC. Right Time, Right Place: Probing the Functions of Organelle Positioning. Trends Cell Biol. 2016;26(2):121–34. Epub 2015/11/07. doi: 10.1016/j.tcb.2015.10.001. PubMed PMID: 26541125.

50. Guardia CM, De Pace R, Sen A, Saric A, Jarnik M, Kolin DA, et al. Reversible association with motor proteins (RAMP): A streptavidin-based method to manipulate organelle positioning. PLoS Biol. 2019;17(5):e3000279. Epub 2019/05/18. doi: 10.1371/journal.pbio.3000279. PubMed PMID: 31100061; PubMed Central PMCID: PMCPMC6542540.

